# Perturbation of protein homeostasis brings plastids at the crossroad between repair and dismantling

**DOI:** 10.1101/2022.07.19.500576

**Authors:** Luca Tadini, Nicolaj Jeran, Guido Domingo, Federico Zambelli, Simona Masiero, Anna Calabritto, Elena Costantini, Sara Forlani, Milena Marsoni, Federica Briani, Candida Vannini, Paolo Pesaresi

## Abstract

The chloroplast proteome is a dynamic mosaic of plastid- and nuclear-encoded proteins. Plastid protein homeostasis is maintained through the balance between *de novo* synthesis and proteolysis. Intracellular communication pathways, including the plastid-to-nucleus signalling and the protein homeostasis machinery, made of stromal chaperones and proteases, shape chloroplast proteome based on developmental and physiological needs. However, the maintenance of fully functional chloroplasts is costly and under specific stress conditions the degradation of damaged chloroplasts is essential to the maintenance of a healthy population of photosynthesising organelles while promoting nutrient redistribution to sink tissues. In this work, we have addressed this complex regulatory chloroplast- quality-control pathway by modulating the expression of two nuclear genes encoding plastid ribosomal proteins PRPS1 and PRPL4. By transcriptomics, proteomics and transmission electron microscopy analyses, we show that the increased expression of *PRPS1* gene leads to chloroplast degradation and early flowering, as an escape strategy from stress. On the contrary, the overaccumulation of PRPL4 protein is kept under control by increasing the amount of plastid chaperones and components of the unfolded protein response (cpUPR) regulatory mechanism. This study advances our understanding of molecular mechanisms underlying chloroplast retrograde communication and provides new insight into cellular responses to impaired plastid protein homeostasis.

## Introduction

Chloroplasts are plant cell organelles of cyanobacterial origin that perform essential metabolic and biosynthetic functions including photosynthesis and fatty acid biosynthesis. The Arabidopsis chloroplast proteome is estimated to consist of several thousand proteins, most of which are encoded by the nuclear genome and post-translationally imported into the organelle (Kmiec *et al*., 2014). The plastid genome, encoding about a hundred proteins, is expressed by transcriptional and translational machineries that conserve many bacteria-like elements. For instance, the chloroplast ribosome shares several features with that of the model organism *E. coli*, used as the reference in early investigations (Mache, 1990). Almost all the chloroplast ribosomal proteins have an orthologue in *E. coli*, however, few differences in ribosome composition, protein domain organisation and function have been described (Bubunenko *et al*., 1994; Yamaguchi *et al*., 2000; Yamaguchi & Subramanian, 2003; Tiller & Bock, 2014; Ahmed *et al*., 2016; Graf *et al*., 2017; Bieri *et al*., 2017; Zoschke & Bock, 2018).

This is the case, for instance, of the S1 protein, encoded by *rpsA* gene, the largest ribosomal protein present in *E. coli* 30S subunit essential for cell viability (Kitakawa & Isono, 1982). S1 promotes the translation initiation step by recognising diverse mRNA leaders and mediating their interaction with the ribosome (Sørensen *et al*., 1998; Hajnsdorf & Boni, 2012). Furthermore, S1 is found in *E. coli* cells both as ribosome-associated- and as free-subunit in cytoplasm (Subramanian, 1983; Kalapos *et al*., 1997; Delvillani *et al*., 2011), and it is responsible for its own post- transcriptional regulation (Skouv *et al*., 1990). S1 bears six homologous non-identical repeats known as S1 domains, members of an ancient RNA binding OB-fold family (Subramanian, 1983; Bycroft *et al*., 1997). It has been proposed that the six S1 domains have a partial functional specialisation, which correlates with their relative position (Salah *et al*., 2009). In particular, domains 1 and 2 mediate the interaction with the ribosome (Giorginis & Subramanian, 1980; Byrgazov *et al*., 2015), domains 3, 4, 5 and 6 are responsible for RNA binding and unfolding (Subramanian, 1983; Duval *et al*., 2013; Cifuentes-Goches *et al*., 2019), with domains 5 and 6 also involved in transcription stimulation and autoregulation (Boni *et al*., 2000; Sukhodolets *et al*., 2006). Recently, domains 4 and 6 have been shown to be implicated in ribosome dimerization and hibernation under stress (Beckert *et al*., 2018).

Unlike S1, the *Arabidopsis thaliana* <u>P</u>lastid <u>R</u>ibosomal <u>P</u>rotein <u>S</u>mall subunit 1 (PRPS1) is characterised by three S1 domains. Its function was characterised by exploiting the knock-down allele *prps1-1*, which produces about one-tenth of wild-type PRPS1 transcripts resulting in one-third of wild-type PRPS1 protein levels in adult plants. Such impairment in *prps1-1* mutants affects the overall plant growth rate and results in pale leaves due to decreased translation in chloroplasts (Romani *et al*., 2012). *A. thaliana* PRPS1 has been found to genetically and physically interact with the nuclear-encoded plastid protein GUN1 (Tadini *et al*., 2016). As a consequence, the depletion of GUN1 in the *prps1-1* genetic background leads to the partial rescue of the mutant phenotype, restoring to wild-type-like levels both PRPS1 abundance and the chloroplast translation capacity (Tadini *et al*., 2016). These results indicate a direct negative regulation of PRPS1, and therefore of chloroplast translation, by GUN1, possibly due to its relations with the chloroplast protein homeostasis machinery (Colombo *et al*., 2016; Tadini *et al*., 2016). Further investigations revealed the involvement of PRPS1 and plastid translation in retrograde signalling upon heat-stress. Under such conditions, the diminished chloroplast translational capacity of *prps1-1* prevents the up- regulation of the *HSFA2* gene, a master regulator of the chloroplast Unfolded Protein Response (Nishizawa *et al*., 2006; Yu *et al*., 2012). Accordingly, seedlings and adult *prps1-1* plants are unable to cope with high temperature showing low survival rates with respect to the wild-type (Yu *et al*., 2012). Interestingly, the introgression of *gun1* mutation in *prps1-1* genetic background rescues its low survival rate in such conditions (Tadini *et al*., 2016). In addition, attempts to constitutively overexpress *PRPS1* gene resulted in a virescent phenotype and in the decrement of PRPS1 protein levels (Yu *et al*., 2012).

Taken together, these observations indicate that chloroplast gene expression is largely controlled at the translational and post-translational level and set the chloroplast translation as a crucial step for the genesis of chloroplast-to-nucleus retrograde signalling pathways (Zoschke & Bock, 2018; Wu *et al*., 2019). Furthermore, the peculiar features of ribosomal protein S1 identified during studies on *E. coli* and *Arabidopsis thaliana*, make it an interesting subject for deeper investigations regarding its role in chloroplast biogenesis, translational regulation and the interconnection with the protein homeostasis maintenance and retrograde communication.

These aspects have been investigated in the present manuscript, where we demonstrate that the knock-out of *PRPS1* gene is incompatible with chloroplast biogenesis and embryo development. Further, we assessed that PRPS1 is unable to functionally replace S1 in *E. coli* cells, and its overexpression inhibits cell growth, as in the case of the endogenous S1 protein (Briani *et al*., 2008; Delvillani *et al*., 2011). We also show that PRPS1 protein accumulation in chloroplasts is prevented post-translationally by the plastid CLP protease complex, while its constitutive over-expression promotes chloroplast degradation via micro- and macro-autophagy (Woodson, 2022), and induces early flowering. This adaptive response is organised at the very beginning of *PRPS1* transcript over- accumulation, as revealed by the transcriptome profile of short-term induced *PRPS1* expression lines. On the contrary, the over-accumulation of Plastid Ribosomal Protein Large subunit 4 (PRPL4) (Bryant *et al*., 2011; Romani *et al*., 2012), here used as control, is tolerated by chloroplasts and leads to the accumulation of transcripts and proteins, such as chaperons and proteases, that are part of the chloroplast-derived Unfolded Protein Response mechanism (cpUPR; Ramundo & Rochaix, 2014; Ramundo *et al*., 2014; Pérez-Martín *et al*., 2014; Llamas *et al*., 2017).

## Materials and methods

### Bioinformatic analyses

S1 domain sequences have been identified using InterPro online tool (https://www.ebi.ac.uk/interpro/). Multiple sequence alignment was performed with MUSCLE online tool (https://www.ebi.ac.uk/Tools/msa/muscle/) and represented as phylogenetic tree employing PhyML (https://toolkit.tuebingen.mpg.de/tools/phyml) and iTOL (https://itol.embl.de/).

### Plant material and growth conditions

The *PRPS1/prps1-2* heterozygous mutant lines were generated by targeting the first exon of *PRPS1* locus in *Arabidopsis thaliana* wild-type (Col-0) genetic background, using the pDe-CAS9 vector described by Fauser et al. (2014) (guide RNA sequence is listed in Table S1). *PRPS1/prps1-2* heterozygous plants, devoid of CAS9 T-DNA, were selected based on the mutation in *PRPS1* sequence. *prps1-2 pPRPS1::PRPS1* complemented lines were obtained by introgressing *PRPS1* genomic locus, including the *pPRPS1* promoter region, in *PRPS1/prps1-2* heterozygous plants and by selecting *prps1-2* viable plants carrying the *pPRPS1::PRPS1* construct. *oePRPS1* and *oePRPL4* lines were obtained by Agrobacterium-mediated transformation of Arabidopsis Col-0 genetic background with the coding sequences of *PRPS1* and *PRPL4* genes, under the control of CaMV35S promoter (pB2GW7 plasmid; https://gatewayvectors.vib.be/). *indPRPS1* and *indPRPL4* lines were obtained by cloning the *PRPS1* and *PRPL4* coding sequences in the Dexamethasone (DEX)-inducible *pOp/LhG4* system (Samalova *et al*., 2005) and by Agrobacterium transformation of Arabidopsis Col-0 plants. *prps1-1* (SAIL_560_B02), *prpl11-1* (GABI_380H05), *clpc1-1* (SALK_014058C) and *clpd-1* (SALK_110649C) T-DNA lines were described in previous works (Pesaresi *et al*., 2001; Sjögren *et al*., 2004; Romani *et al*., 2012; Pulido *et al*., 2016) and manually crossed for obtaining *prps1-1 prpl11-1, prps1-1 clpc1-1, prps1-1 clpd-1* double mutants. Primers required for gene cloning and mutant line isolation are listed in Table S1. Wild-type and mutant seeds were grown on soil in climate chambers under long-day (150 μmol m^−2^ sec^−1^ 16 h/8 h light/dark cycles) and short-day (150 μmol m^−2^ sec^−1^ 8 h/16 h light/dark cycles) conditions. For growth experiments on Dexamethasone, seeds were surface-sterilised and grown for 16 days (80 μmol m^−2^ sec^−1^ on a 16 h/8 h light/dark cycle) on Murashige and Skoog medium (Duchefa) supplemented with 1% (w/v) sucrose and 1.5% Phyto-Agar (Duchefa), Dexamethasone was added at the final concentration of 2 µM. Growth rate was determined by ImageJ software (https://imagej.nih.gov/).

### Whole-mount preparation and optical microscopy

To analyse defects in embryo development, siliques of Col-0 and heterozygous *PRPS1/prps1-2* plants were manually dissected and cleared as reported in Tadini *et al*., 2018. Developing seeds were observed using a Zeiss Axiophot D1 microscope equipped with differential interface contrast optics. Images were documented with an Axiocam MRc5 camera (Zeiss).

### E. coli strains

*PRPS1* coding sequence was cloned into pQE31-pREP4 plasmid system (primers are listed in Table S1), under the control of bacteriophage T5 promoter fused upstream to *lacO* operator sequences. *PRPS1*-*pQE31-pREP4* and *rpsA-pQE31-pREP4* plasmids (Briani *et al*., 2008) were then transferred into *araBp-rpsA* conditional expression strain C-5699, in which the *rpsA* gene is expressed in presence of 1% (w/v) arabinose and repressed in presence of 0.4% (w/v) glucose. To obtain the *PRPS1* overexpressing strain, *PRPS1*-*pQE31-pREP4* plasmids were introduced into the *E. coli* C-1a strain (Sasaki & Bertani, 1965), while *rpsA-pQE31-pREP4* strain C-5691 was used as control (Briani *et al*., 2008; Delvillani *et al*., 2011).

### Chlorophyll a fluorescence measurements

The Imaging Chl a fluorometer (Walz Imaging PAM; https://www.walz.com/) was used to determine Chl fluorescence *in vivo*. Eight plants for each genotype and condition were analysed at 18 days after sowing (DAS). Average values plus-minus standard deviations were then calculated. 20 min dark- adapted plants were exposed to blue measuring light (intensity 4) and a saturating light flash (intensity 4) was used to calculate the maximum quantum yield of PSII, *Fv/Fm*.

### Nucleic acid analyses

For qRT-PCR analyses, 1 µg of total RNA were treated with iScript™ gDNA Clear cDNA Synthesis Kit (Bio-Rad; https://www.bio-rad.com/) for genomic DNA digestion and first-strand cDNA synthesis. qRT-PCR analyses were performed on a CFX96 Real-Time system (Bio-Rad; https://www.bio-rad.com/), using primer pairs listed in Table S1. *PP2AA3* (*AT1G13320*) transcripts were used as internal reference, as described in Czechowski et al. (2005). Data obtained from three biological and three technical replicates for each sample were analysed with the Bio-Rad CFX Maestro 1.1 (v 4.1) (Bio-Rad; https://www.bio-rad.com/).

### *In vivo* Translation Assay

The *in vivo* translation assay was performed essentially as previously described (Tadini *et al*., 2012). *indPRPS1* leaf discs (6 mm in diameter) were vacuum-infiltrated in liquid MS medium supplemented with 1% (w/v) sucrose and, where indicated, 2 µM Dexamethasone. After 6-hours exposure to 80 μmol photons m^−2^ s^−1^ white light, leaf discs were incubated with a buffer containing 1 mM K_2_HPO_4_– KH_2_PO_4_(pH 6.3) and 0.05% (v/v) Tween-20, supplemented with 20 μg/ml cycloheximide, to inhibit cytosolic translation. [^35^S]methionine was then added (0.1 mCi/ml) and leaf discs were vacuum- infiltrated and exposed to light (80 μmol photons m^−2^ s^−1^). 5 leaf discs were collected at each time point (15 and 30 min). Total proteins extraction and Tris-glycine SDS-PAGE fractionation is described below. Signals were detected using the Phosphorimager GE Healthcare Life Sciences (https://www.gehealthcare.com/).

### Isolation of PRPS1-containing protein complexes

The isolation of PRPS1-containing complexes was performed according to previous works (Barkan, 1998; Merendino *et al*., 2003). 100 mg of leaf fresh weight were ground in liquid nitrogen and resuspended in 1 ml 0.2 M Tris-HCl, pH 9, 0.2 M KC1, 35 mM MgC1_2_, 25 mM EGTA, 0.2 M sucrose, 1% Triton X-100, 2% polyoxyethylene-10-tridecyl ether, supplemented with 500 µg/ml heparin, 100 µg/ml chloramphenicol and 25 µg/ml cycloheximide. The extract was then solubilised with 0.5% (w/v) sodium deoxycholate for 5 min on ice. After centrifugation (15 min at 10000g), 800 µl of supernatant was loaded onto 3.6 ml 15-55% (w/v) sucrose gradients in polysome gradient buffer (40 mM TrisHCl pH 8, 20 mM KCl, 10 mM MgCl_2_, 100 µg/ml chloramphenicol and 500 µg/ml heparin). Sucrose gradients were centrifuged in SW60 rotors (Beckman) for 18 h at 180000 g at 4 °C. 9 fractions of 400 µl each were collected from the top of the tube and subjected to SDS-PAGE fractionation. *E. coli* ribosome fractionation was performed accordingly to Delvillani et al. (2011).

### Transmission electron microscopy (TEM)

TEM analyses were performed as described previously (Jeran *et al*., 2021). Plants were grown for 16 days on MS synthetic medium supplemented with 1% (w/v) sucrose and, where indicated, 4 µM Dexamethasone. Plant material was vacuum-infiltrated with 2.5% glutaraldehyde, in 100 mM sodium cacodylate buffer, for 4 h at room temperature and incubated overnight at 4 °C. Samples were rinsed twice with 100 mM sodium cacodylate buffer for 10 min each, and post-fixed in 1% (w/v) osmium tetroxide in 100 mM cacodylate buffer for 2 h at 4 °C. After washings, samples were counterstained with 0.5% (w/v) uranyl acetate overnight at 4 °C, in the dark. The tissues were then dehydrated by increasing concentrations of ethanol (70%, 80%, 90%; v/v), 10 min each. Samples were then dehydrated with 100% ethanol for 15 min and permeated twice with 100% propylene oxide for 15 min. Epon-Araldite resin was prepared mixing properly Embed-812, Araldite 502, dodecenylsuccinic anhydride (DDSA) and Epon Accelerator DMP-30. Samples were infiltrated first with a 1:2 mixture of Epon-Araldite and propylene oxide for 2 h, then with Epon-Araldite and propylene oxide (1:1) for 1 h and left in a 2:1 mixture of Epon-Araldite and propylene oxide overnight at room temperature. Samples were then incubated in pure resin before polymerisation at 60 °C for 48 h. Ultra-thin sections of 70 nm were then cut with a diamond knife (Ultra 45°, DIATOME) and collected on copper grids (G300-Cu, Electron Microscopy Sciences). Samples were observed by transmission electron microscopy (Talos L120C, Thermo Fisher Scientific) at 120 kV. Images were acquired with a digital camera (Ceta CMOS Camera, Thermo Fisher Scientific).

### Protein sample preparation and immunoblot analyses

For immunoblot analyses, total proteins were prepared as described (Tadini *et al*., 2020c). Plant material was homogenised in Laemmli sample buffer [20% (v/v) glycerol, 4% (w/v) SDS, 160 mm Tris–HCl pH 6.8, 10% (v/v) 2-mercaptoethanol] to a concentration of 0.1 mg µl^−1^ (leaf fresh weight/Laemmli sample buffer). Samples were incubated for 15 min at 65°C and, after a centrifugation step (10 min at 16 000 g), the supernatant was incubated for 5 min at 95 °C. Protein samples corresponding to 4 mg (fresh weight) of seedlings were fractionated by SDS–PAGE 10% (w/v) acrylamide (Schägger & von Jagow, 1987) and then transferred to polyvinylidene difluoride (PVDF) membranes (0.45 µm pore size). Replicate filters were immunodecorated with specific antibodies. Antibodies directed against AtHsp90-1 (AS08 346) and ClpB3 (AS09 459) were obtained from Agrisera (https://www.agrisera.com/), AtHsc70-4 antibody was obtained from Antibodies- online (https://www.antibodies-online.com/), antibodies directed against plastid ribosomal proteins (PRPS1, PRPL4 and PRPS5) were obtained from Uniplastomic, while polyclonal S1 antibody was kindly donated by U. Bläsi (University of Vienna).

### Transcriptome analysis

Total RNA was extracted from leaf discs harvested from *indPRPS1* and *indPRPL4* plants and vacuum infiltrated in either the absence or presence of DEX for 6 hours for a total of 5 biological replicates for each group. Total RNA extraction was performed using RNeasy Mini Kit (Qiagen), according to the manufacturer’s instructions. RNA concentrations and integrity were determined via NanoDrop One C (ThermoFisher Scientific) and agarose-gel electrophoresis. Extracted RNA samples were sent to Novogene for sequencing via high throughput Illumina NovaSeq platform which employ a paired- end 150 bp sequencing strategy. Raw data were processed and mapped to the Arabidopsis genome TAIR10 using STAR-RSEM software (Li & Dewey, 2011). Differentially Expressed Genes (DEGs) were identified through the R package EdgeR (v 3.15; Robinson *et al*., 2009). Called DEGs were statistically filtered via Benjamini-Hochberg False Discovery Rate method (FDR < 0.05). GO enrich- ment analyses were performed using agriGO v2.0 online tool and further processed by REVIGO (Supek *et al*., 2011; Tian *et al*., 2017). The RNA-seq data were deposited in the Gene Expression Omnibus data repository under the dataset identifier GSE205271.

### Proteome analysis

Proteins were extracted from 1 g of plantlets following SDS/phenol method as described in Vannini et al. (2021). Proteins were then digested with trypsin via Filter Aided Sample Preparation (FASP) (Wiśniewski, 2019). Peptides were analysed by LC-MS/MS as described by Paradiso et al. (2020). Briefly, after LC separation peptides were sprayed into the mass spectrometer and eluting ions were measured in an Orbitrap mass analyser set at a resolution of 35000 and scanned between m/z 380 and 1500. Data dependent scans (top 20) were employed to automatically isolate and generate fragment ions by higher energy collisional dissociation (HCD); Normalised collision energy (NCE): 25% in the HCD collision cell and measurement of the resulting fragment ions was performed in the Orbitrap analyser, set at a resolution of 17500. Peptide ions with charge states of 2^+^ and above were selected for fragmentation. Raw data were searched against the *Arabidopsis thaliana* TAIR protein database (2010 version) with MaxQuant program (v.1.5.3.3), using default parameters. For the quantitative analysis, the “ProteinGroups” output files were filtered to retain only protein groups detected with at least two peptides in at least three of the four biological replicates, and in at least one analytical group. The mass spectrometry proteomics data have been deposited in the ProteomeXchange Consortium via the PRIDE (Perez-Riverol *et al*., 2019) partner repository under the dataset identifier PXD034479. Missing values were replaced with the R package imputeLCMD (v.2.1) using Hybrid imputation method: imputation of left-censored missing data (missing values ≥ 50% of number of replicas) was done using QRILC method, instead missed at random data (< 50% of replicas) were imputed using KNN method. Log2 transformed LFQ intensities were centred by Zscore normalisation method of Perseus (https://www.maxquant.org/perseus/) and then subjected to Student’s Tests (S0=0.1, FDR<0.05) in order to discover Differentially Abundant Proteins (DAPs). Hierarchical clustering analysis was carried out using Perseus software and default parameters. DAPs categorisation was achieved using TAIR GO annotation tool (https://www.arabidopsis.org/tools/bulk/go/index.jsp). GO term enrichment analysis was performed using the PANTHER classification system (Mi et al., 2021; http://geneontology.org).

## Results

### Depletion of *PRPS1* gene leads to embryo lethality

Previous studies reported on the Arabidopsis *prps1-1* knock-down mutant phenotype, characterised by pale-green leaves, reduced growth rate and photosynthetic performance, as result of hampered plastid protein synthesis (Romani *et al*., 2012; Yu *et al*., 2012). However, the consequences of the complete inhibition of PRPS1 protein accumulation in Arabidopsis plastids has not yet been investigated. To fill this gap, the nuclear *PRPS1* gene was edited by targeting the first exon of *PRPS1* coding sequence using the CRISPR/Cas9 technology. Plant lines devoid of the Cas9 gene were obtained in T2 generation and the DNA region complementary to the designed guide RNA was sequenced (primers and guide RNA sequences are listed in Table S1). Only *PRPS1/prps1* heterozygous plants, with wild-type-like plant size, leaf pigmentation and photosynthetic performance, could be isolated and the resulting *prps1-2* allele showed the deletion of a Cytosine in the first exon (+80 bp from the transcription starting site), right downstream the ATG translation start codon (+ 4 bp from ATG, Fig.1 A, B). This event disrupts the *PRPS1* reading frame and introduces a premature Umber STOP codon in place of Leu-11 (+108 bp from the transcription starting site) (Fig. 1A). Furthermore, only *PRPS1/prps1-2* heterozygous and *PRPS1/PRPS1* homozygous plants could be identified within the progeny of the self-fertilised *PRPS1/prps1-2* heterozygous line, showing the 2-to-1 mendelian segregation ratio typical of mutations causing embryo lethality, as in the case of other plastid ribosomal protein knock-out mutants (Bryant *et al*., 2011; Romani *et al*., 2012; Yin *et al*., 2012). Accordingly, the observation of *PRPS1/prps1-2* developing siliques at 10 Days After Fertilisation (DAF) revealed one-quarter of the seeds to be albino (Fig.1 C), indicating that *PRPS1* is essential during early stages of embryogenesis and seed development. In particular, optical section of cleared, whole-mount seeds from *PRPS1/prps1-2* siliques at 3 DAF showed that around 25% of the embryos were arrested at the globular development stage, displaying a disorganised cell division pattern similar to the ones previously described for other knock-out mutants in essential plastid ribosomal proteins (Romani et al., 2012; Fig. 1 D). Furthermore, the defect in embryo development was fully rescued in *prps1-2 pPRPS1::PRPS1* plants, obtained by introducing the *PRPS1* genomic DNA into the *PRPS1/prps1-2* genetic background and isolating wild-type homozygous *prps1-2* plants carrying the *pPRPS1::PRPS1* construct (Fig. 1 C, D).

**Figure 1.**
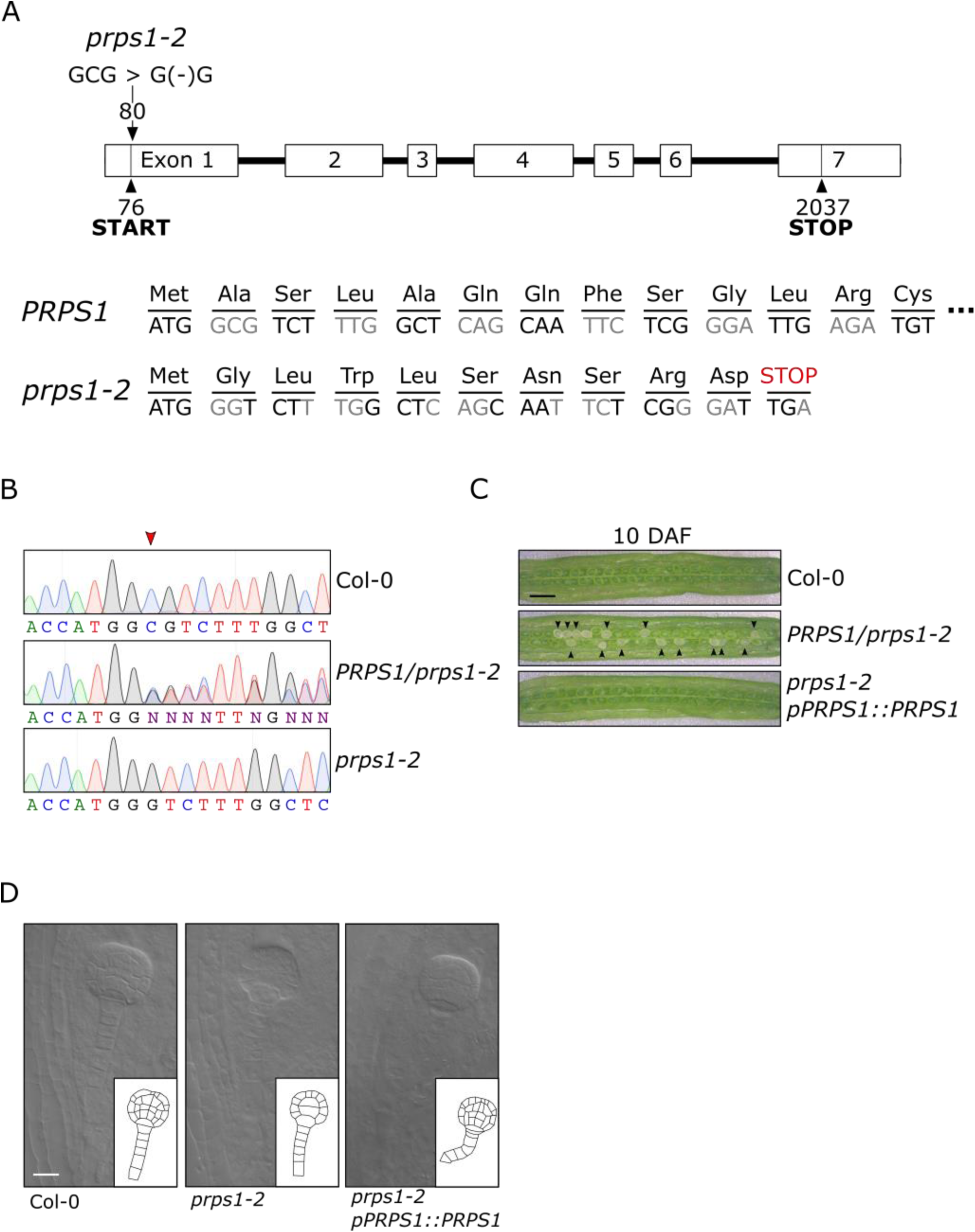
Molecular description of *prps1-2* allele and the corresponding embryo-lethal phenotype. A) Schematic representation of *PRPS1* gene. Exons are indicated as numbered white boxes, and introns as black lines. Positions of start and stop codons with respect to the transcription initiation site, as well as the CRISPR/Cas9-induced deletion (*prps1-2*), are indicated. The *PRPS1* reading frame and corresponding amino acids in wild-type and *prps1-2* mutant are also reported. B) Sequencing electropherogram showing the single nucleotide deletion (coding strand) in *prps1-2* hetero- and homozygous mutants with respect to Col-0. The red arrowhead indicates the position of the CRISPR/Cas9-induced deletion. “N” readings in *PRPS1/prps1-2* lines results from double peaks, generated by Cas9-induced deletion in one of the two homologous chromosomes. C) Morphological characteristics of developing seeds in siliques at 10 Days After Fertilization (DAF) of Col-0, heterozygous *PRPS1/prps1-2* and *prps1-2* complemented with *pPRPS1::PRPS1* genomic locus. Around one quarter of white seeds are clearly distinguishable among the green seeds of *PRPS1/prps1- 2* siliques. Bars = 2 mm. D) Cleared whole mount of Col-0, heterozygous *PRPS1/prps1-2* and *prps1- 2 pPRPS1::PRPS1* complemented seeds containing embryos at the globular stage (3 DAF). Bars = 20 µm.

### *PRPS1* over-expression impairs chloroplast activity and biogenesis

Besides the embryo lethal phenotype caused by the *prps1-2* allele and the slightly pale cotyledons and true leaves, together with hampered photosynthetic efficiency (*F_V_/F_M_*), typical of plants carrying the knock-down *prps1-1* allele (Fig. 2 A; see also Romani et al., 2012; Yu et al., 2012), the *CaMV35S*- mediated over-expression of *PRPS1* gene in the Arabidopsis Col-0 background (*oePRPS1*), also resulted in a visible phenotype, characterised by virescent young leaves with a marked drop in *F_V_/F_M_*values (Fig. 2 A). Interestingly, *oePRPS1* seedlings were incapable of over-accumulating the PRPS1 protein (Fig. 2 B), showing an accumulation level lower than the one observed in *prps1-1* leaves, despite the *PRPS1* transcript level having been about two-fold the Col-0 control leaves (Fig. 2 C, see also Tadini et al. 2016). On the contrary, Col-0 plants carrying the *CaMV35S::PRPL4* construct (*oePRPL4*), here used as the control, were able to accumulate up to 25-30 fold more *PRPL4* transcripts and almost double the amount of PRPL4 protein without affecting chloroplast biogenesis and activity, as shown by *oePRPL4* lines indistinguishable from Col-0 (Fig. 2 A, B, C).

**Figure 2.**
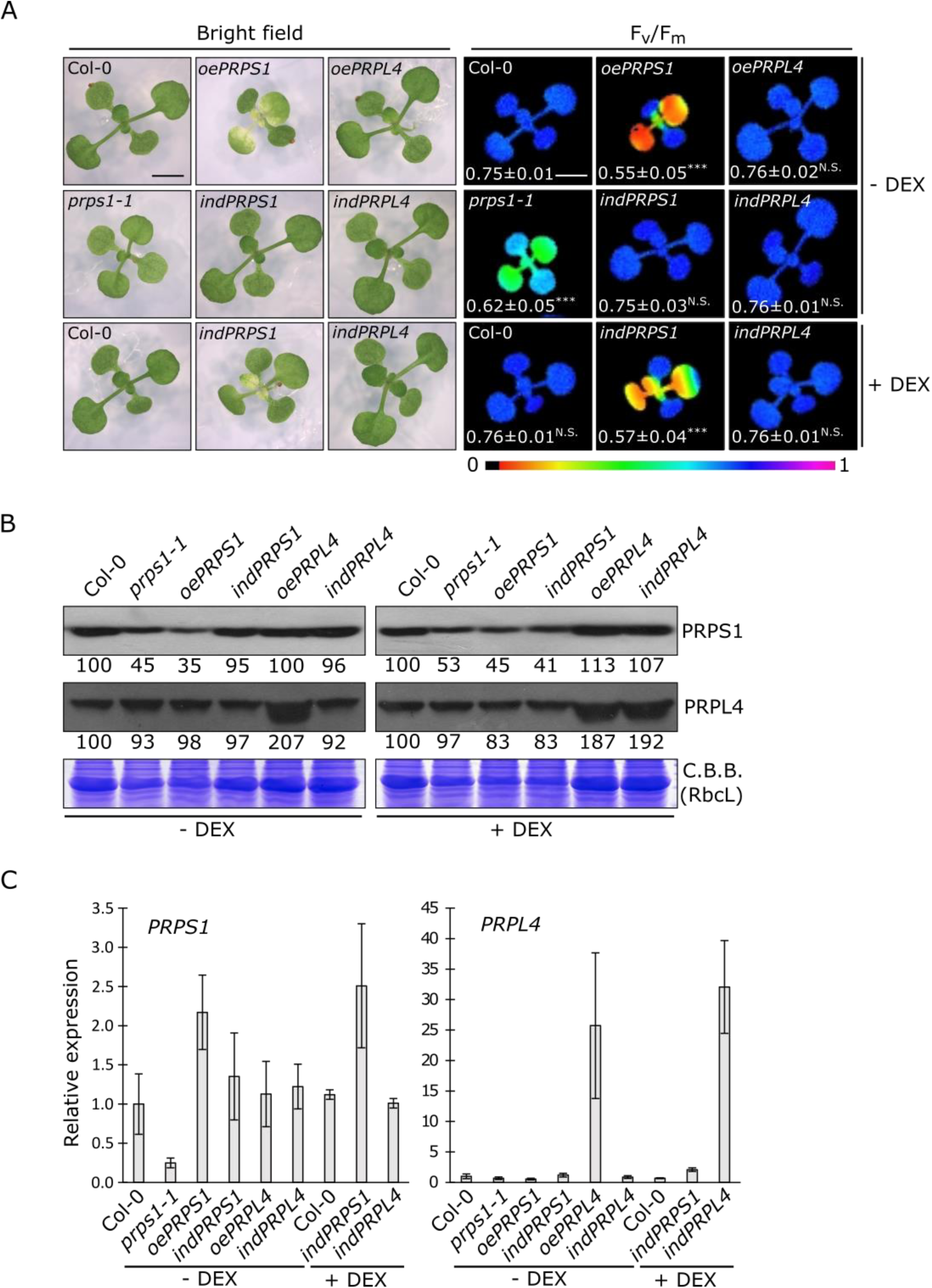
Visible and molecular phenotypes of plantlets carrying altered amount of PRPS1 and PRPL4 proteins. A) Visible phenotypes and photosynthetic efficiency (*F_V_/F_M_*) of 16 Days After Sowing (DAS) plantlets grown on MS medium devoid (- DEX) or supplemented (+ DEX) with 2 µM dexamethasone. On the left (bright field), the visible phenotypes shown by Col-0, the knock-down *prps1-1* mutant, *oePRPS1* and *oePRPL4* constitutive over-expressing lines, *indPRPL4* and *indPRPS1* inducible lines. On the right panel (*F_V_/F_M_*), the photosynthetic efficiency of each genotype is shown both as false colour imaging and average values ± standard deviations. Statistical significance calculated via Student’s t-test (*** indicates *P* < 0.001; N.S. not significant). Scale bar = 1 cm. B) Immunoblots of total protein extracts from Col-0, *prps1-1*, *oePRSP1*, *indPRPS1*, *oePRPL4* and *indPRPL4* 16 DAS plants grown on MS medium devoid (- DEX) or supplemented (+ DEX) with 2 µM dexamethasone. PRPS1 and PRPL4 specific antibodies were used for immuno-decoration. Coomassie Brilliant Blue (C.B.B.) stained gels are shown as loading control. Numbers show the relative protein abundance with respect to Col-0 (indicated as 100). Standard deviation was below ± 15%. One filter out of three biological replicates is shown. C) Relative expression values of *PRPS1* and *PRPL4* genes determined by qRT-PCR analyses of total RNA extracted from Col-0, *prps1-1*, *oePRSP1*, *indPRPS1*, *oePRPL4* and *indPRPL4* 16 DAS plants grown on MS medium devoid (- DEX) or supplemented (+ DEX) with 2 µM dexamethasone. Results of one out of three biological replicates are shown. Error bars indicate standard deviations of three technical replicates.

In order to investigate this aspect further, *PRPS1* and *PRPL4* coding sequences were cloned into the *pOp/LhG4* vector, which allows the inducible over-expression of the two genes once the glucocorticoid analogue dexamethasone (DEX) is provided. The two constructs were introduced into Arabidopsis Col-0 genetic background, via Agrobacterium-mediated transformation, resulting in the dexamethasone-inducible lines *indPRPS1* and *indPRPL4*. In the absence of DEX, the *indPRPS1* line was virtually indistinguishable from Col-0 when grown on MS medium under sterile conditions for 16 days (Fig 2 A). Conversely, when the growth medium was supplemented with 2µM DEX, *indPRPS1* seedlings showed a leaf virescent phenotype and a drop in photosynthetic performance, together with a reduced accumulation of PRPS1 protein resembling the phenotype of *prps1-1* and *oePRPS1* lines (Fig 2 A, B). This was despite *PRPS1* transcripts accumulation to levels higher than Col-0 control leaves (Fig. 2 C). On the other hand, the inducible overexpression of *PRPL4* resulted in an increased accumulation of *PRPL4* transcripts and protein, similar to *oePRPL4* seedlings (Fig. 2 B, C), without any impact on chloroplast activity and leaf greening (Fig. 2 A).

To understand how chloroplasts ultrastructure organization is affected by the increased expression of *PRPS1* gene, thin sections of emerging young leaves from 16 DAS seedlings were observed under Transmission Electron Microscopy (TEM; Fig. 3). As expected, Col-0, *prps1-1, indPRPS1* - DEX, *indPRPL4* ± DEX, and *oePRPL4* mesophyll cells displayed properly developed chloroplasts with the typical organization in grana stacks and stroma lamellae (Fig. 3). However, both the induction (*indPRPS1* + DEX) and the constitutive increased expression of *PRPS1* caused the formation of miss-shaped and swollen chloroplasts containing enlarged plastoglobuli in the stroma (Fig. 3). Furthermore, large budding vesicles with electron dense material were detectable, suggesting ongoing chloroplast degradation, resembling the fission-type ATG-independent micro-autophagy (Woodson, 2016, 2022; Tadini *et al*., 2020c; Jeran *et al*., 2021). In some cases, entire round-shaped chloroplasts, detached from the plasma membranes and with still recognizable grana stacks, were observed inside the vacuole, compatible with the ATG-dependent micro-autophagy process (Fig. 3J; Zhuang and Jiang, 2019; Woodson, 2022). To further prove that chloroplasts are indeed undergoing vacuole-mediated degradation, we investigated the relative expression of genes associated with either the chloroplast ATG-dependent or ATG-independent chloroplast quality-check and degradation pathways (Fig. S1). Strikingly, both *ATI1* (Michaeli *et al*., 2014) and *ATG8f* (Liu *et al*., 2021) transcripts were highly enriched in plantlets with increased *PRPS1* transcript accumulation, driven by either *CaMV35S-* or DEX-induced promoters (Fig. S1 A), whereas *NPC1* and *VPS15* (Lemke *et al*., 2021) were the only genes of the ATG-independent pathway significantly up-regulated in *oePRPS1* plantlets (Fig. S1 B).

**Figure 3.**
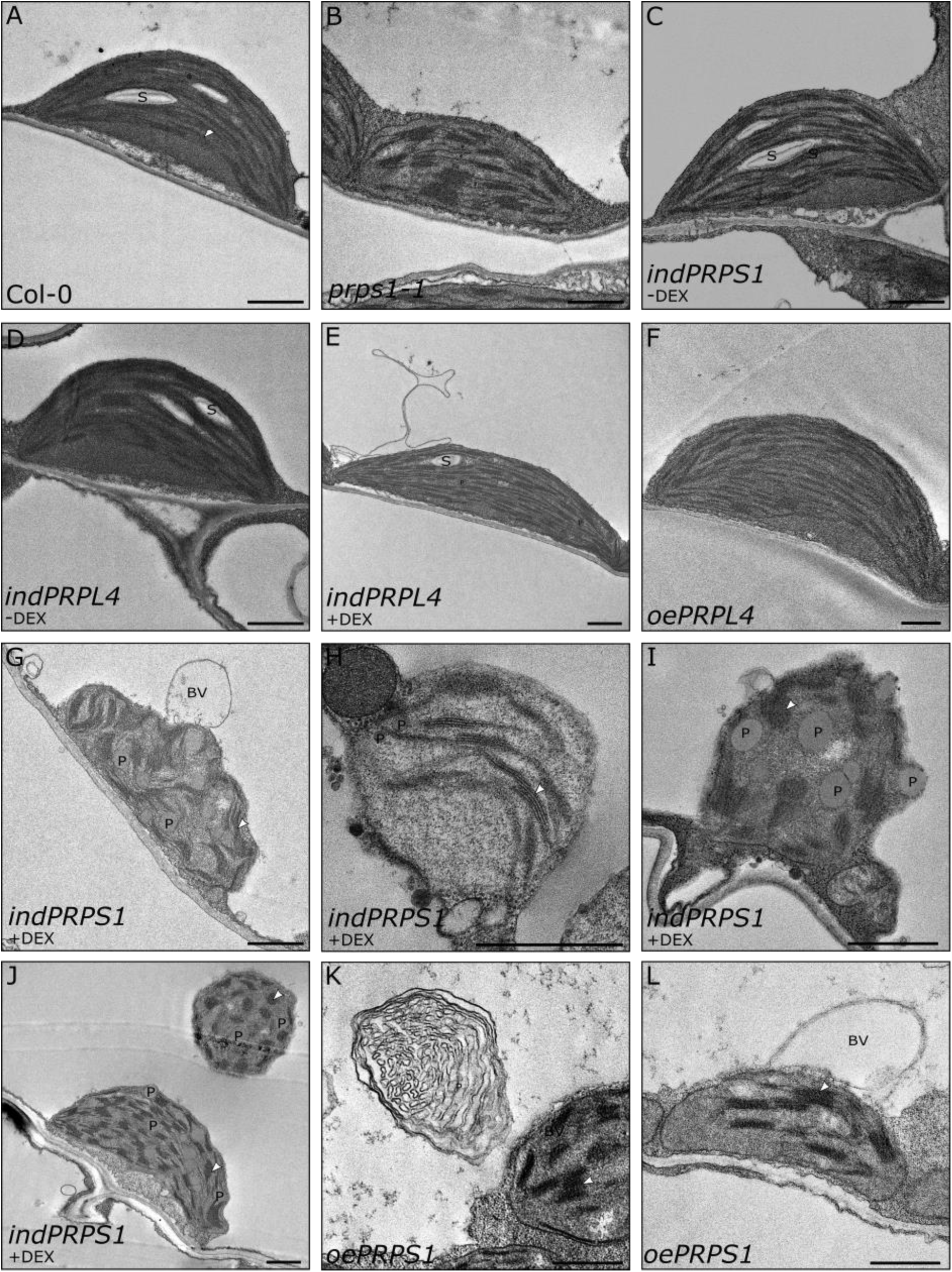
-. TEM micrographs of chloroplasts in mesophyll cells of Col-0 (A) control, *prps1-1* (B), *indPRPS1* (C, G, H, I, J), *indPRPL4* (D, E), *oePRPL4* (F) and *oePRPS1* (K, L) leaves obtained from 16 DAS plantlets grown on MS medium supplemented, or not, with DEX. The young portion of leaves, proximal to the petiole, showing the yellow to pale-green phenotype in *oePRPS1* and *indPRPS1* lines, was used for the analyses. S: starch granules; White arrowhead: thylakoid membranes; BV budding vesicles; P: Plastoglobules.

### *PRPS1* short-term increased expression inhibits plastid protein translation

To gain a dynamic view on PRPS1 function, the kinetics of *PRPS1* transcript and protein accumulation was monitored in leaf discs (6 mm in diameter) from young leaves of *indPRPS1* plantlets infiltrated with 2 µM DEX. Leaf discs were sampled at 0, 3, 6 and 24 hours after infiltration (HAI) and transcript and protein accumulation were monitored by RT-qPCR (Fig. 4 A) and immunoblotting (Fig. 4 B), respectively. As control, the same experimental set-up was used to monitor the induction of *PRPL4* expression (Fig. 4 A, C). The accumulation of *PRPL4* and *PRPS1* mRNAs indicated that both inducible lines were able to specifically express the related genes with comparable kinetics and transcript amounts (Fig. 4 A). Both lines reacted to the presence of DEX showing a high expression level of the related transcripts at 3 HAI, reaching the peak at 6 HAI and a significant decrease at 24 HAI. Nevertheless, the induction of *PRPS1* expression failed to yield the over-accumulation of PRPS1 protein. Indeed, PRPS1 protein level remained stable until 3 HAI, while diminished to almost undetectable levels from 6 to 24 HAI (Fig. 4 B). Interestingly, other plastid ribosomal proteins, such as PRPL4 and PRPS5, showed a marked decreased over time, indicating a general alteration of plastid ribosome accumulation, and possibly of plastid translation. On the contrary, the 24-hour induction of *PRPL4* resulted in more than two-fold accumulation of PRPL4 protein with respect to time 0 (Fig. 4 C). In particular, the increase in PRPL4 protein accumulation was observed over time, starting from 3 HAI and reaching the largest amount at 6 HAI, while the accumulation of PRPS1 and PRPS5 plastid ribosomal proteins remained unaltered.

**Figure 4.**
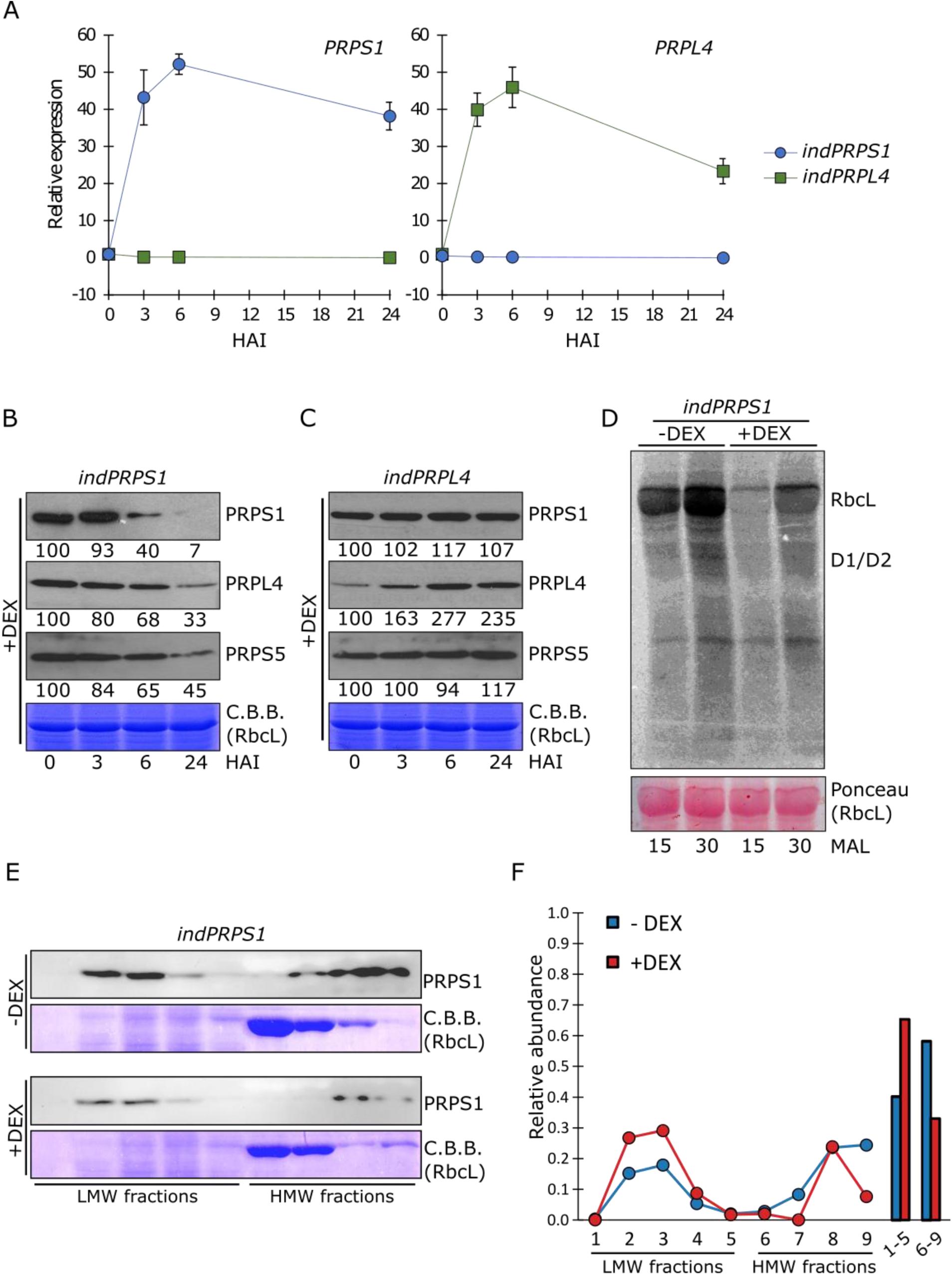
Molecular features of *indPRPS1* and *indPRPL4* lines upon DEX-induced gene expression. A) Relative expression values of *PRPS1* and *PRPL4* genes determined by qRT-PCR analyses of total RNA extracted from *indPRPS1* (circles, blue) and *indPRPL4* (squares, green) leaf discs sampled at 0, 3, 6 and 24 Hours After Infiltration (HAI) with 2 µM dexamethasone (DEX). Data from one out of three biological replicates are shown. Error bars indicate standard deviations of three technical replicates. B) Immunoblots of total protein extracts from *indPRPS1* leaf discs sampled at 0, 3, 6 and 24 HAI with 2 µM DEX. Filters were incubated with antibodies raised against PRPS1, PRPL4 and PRPS5 plastid ribosomal proteins. C.B.B.-stained gels are shown as loading control. Numbers show the relative protein abundance with respect to 0 HAI (indicated as 100). Data from one out of three biological replicates are shown. Standard deviation was below 15%. C) Immunoblots of total protein extracts from *indPRPS4* leaf discs, sampled and analysed as in B, used as control. D) Total protein extracts from Col-0 and *indPRPS1* leaf discs previously incubated in DEX-containing MS medium for 6 hours and then infiltrated with ^35^S-Methionine in the presence of cycloheximide, an inhibitor of cytosolic protein synthesis. The leaf proteins were sampled 15 and 30 minutes after labelling (MAL). RbcL and D1/D2 proteins signals are indicated. The ponceau-stained filter is shown as loading control. E) *indPRPS1* ± DEX (6 HAI) leaf material was subjected to sucrose gradient fractionation aimed to isolate PRPS1-contaning complexes. Filters were immuno-decorated with PRPS1 antibody to show the accumulation of PRPS1 in Low Molecular Weight (LMW) and High Molecular Weight (HMW) fractions. C.B.B.-stained gels are shown as loading control. F) Quantification of PRPS1 accumulation in LMW (1-5) and HMW (6-9) fractions as evaluated by Image Lab software on representative blots.

To investigate the possible negative effect of *PRPS1* inducible expression on plastid protein translation, *indPRPS1* leaf discs were incubated in MS medium (± DEX) for 6 hours and then infiltrated with ^35^S-Methionine and cycloheximide, allowing for the detection of *de novo* synthesized plastid-encoded proteins, while blocking the cytosolic translation. As shown by the pulse-labelling experiment, the synthesis rate of RbcL and D1/D2 proteins were markedly reduced in *indPRPS1* + DEX, over 15 and 30 minutes, when compared to *indPRPS1* leaf discs in the absence of dexamethasone (- DEX; Fig. 4 D), proving that *PRPS1* inducible over-expression leads to plastid translation inhibition. This aspect was investigated further by isolating the PRPS1-containing complexes and monitoring the PRPS1-to-ribosome stoichiometry and the accumulation of “free” PRPS1 fraction, as previously reported in *E. coli* (Delvillani *et al*., 2011). To do so, *indPRPS1* leaf material (6 hours ± DEX) was subjected to sucrose gradient fractionation and probed with the PRPS1 antibody (Fig. 4 E). As observed in Chlamydomonas (Merendino *et al*., 2003), PRPS1 protein was found in two distinct populations, as “Low Molecular Weight (LMW, free PRPS1 fraction)”, bound solely to the mRNA, and as “High Molecular Weight (HMW)”, corresponding to the S1 fraction bound to the ribosome core. The increase of PRPS1 presence in the LMW fractions of *indPRPS1* + DEX line, compared to the - DEX counterpart (Fig. 4 E, F), supports further the inhibition of plastid translation observed in Fig. 4 D. Consistent with these data, we observed that PRSP1 was almost absent in HMW fraction in *oePRPS1* samples (Fig. S2).

### The over-accumulation of Arabidopsis PRPS1 protein inhibits *Escherichia coli* cell growth

Similarly to Arabidopsis, the depletion of the ribosomal protein S1 (Rps1) in *E. coli* cells leads to lethality (Kitakawa & Isono, 1982). Moreover, the down-regulation as well as the over-accumulation of Rps1 impairs protein translation by altering the stoichiometry between Rps1 and the ribosome core, leading to bacteriostatic effects (Briani et al., 2008; Delvillani et al., 2011; see also Fig. 5). The *E. coli* S1 protein is 557 aa long with a molecular weight of 61.2 kDa and possesses six S1 domains (Bycroft et al., 1997; Fig. 5 A) that are typical of several RNA binding proteins (Murzin, 1993; Arcus, 2002; Theobald *et al*., 2003). In *A. thaliana*, the S1 homologous PRPS1 is markedly smaller, with 373 aa and a molecular weight of 40.5 kDa, as mature form. The *in silico* analysis of PRPS1 amino acid sequence identified three S1 domains (Fig. 5 A), in agreement with early analyses of spinach S1 protein (Franzetti *et al*., 1992). In addition, PRPS1 protein shows a high degree of identity (*i.e.* about 50%) with the S1 ribosomal proteins from cyanobacteria, which possess three S1 domains as well (Sugita *et al*., 1995; Salah *et al*., 2009). In order to investigate the possible relationships between the three S1 domains identified in PRPS1 and the six S1 domains in *E. coli* S1, the amino acidic sequences of each S1 domain were aligned and clustered in a phylogenetic tree (Fig. S3). The resulting tree showed that domains 1 and 2 of PRPS1 are more similar to the corresponding domains of S1, while the PRPS1 domain 3 clusters together with the domains 3, 4 and 5 of S1.

**Figure 5.**
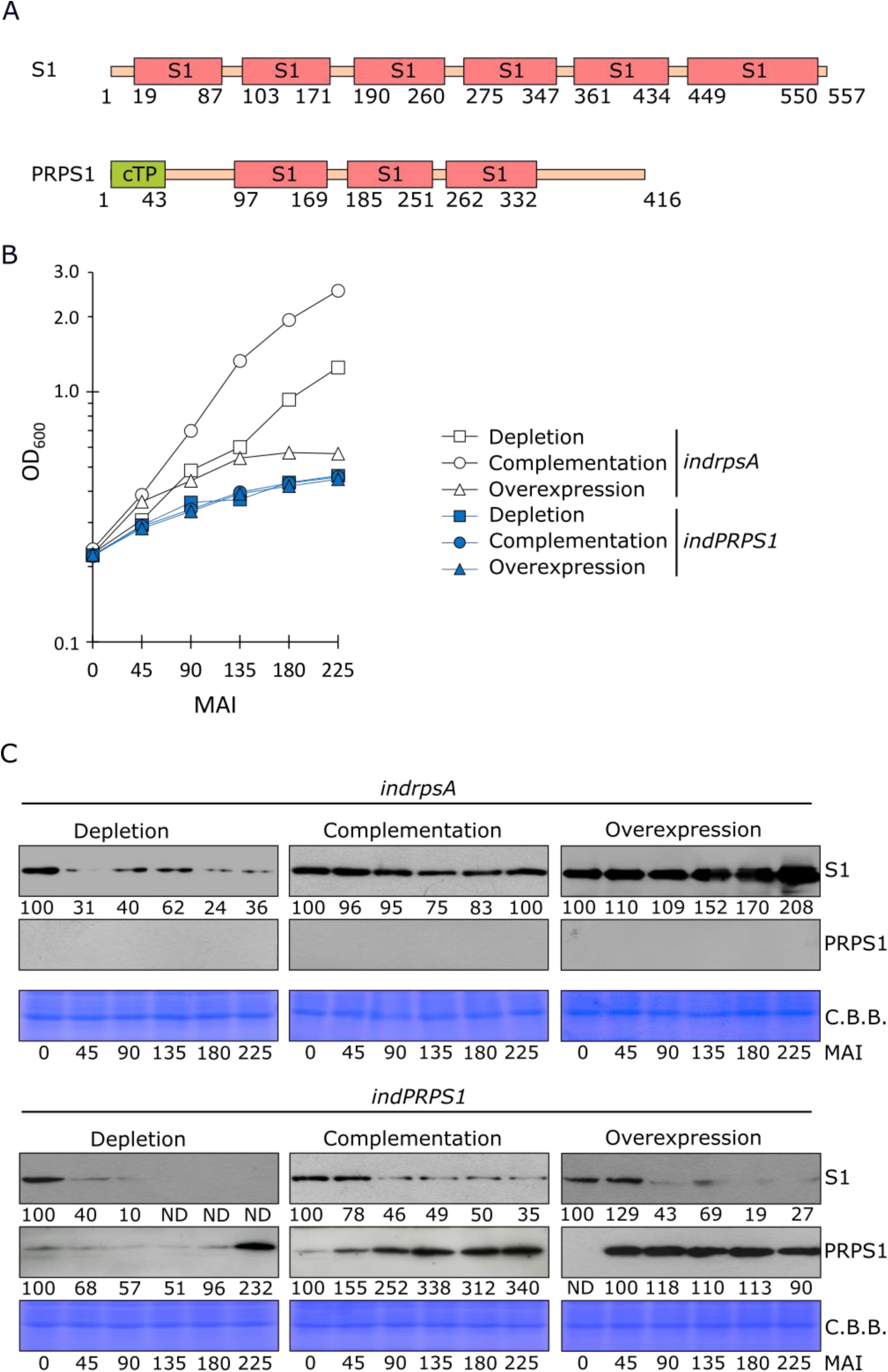
Comparison of PRPS1 and S1 protein activities in *E. coli* cells. A) Schematic representation of S1 and PRPS1 proteins. S1 domains are depicted as pink boxes, while the chloroplast transit peptide (cTP) of PRPS1 protein as green box. Numbers indicate the position of amino acid residues. Domain predictions are based on InterPro online tool (https://www.ebi.ac.uk/interpro/). B) OD_600_measurements of *E. coli indrpsA* (white symbols) and *indPRPS1* (blue symbols) strains grown under Depletion (0.4% glucose; squares), Complementation (0.4% glucose + 0.01 mM IPTG; triangles) or Overexpression (1% arabinose + 1 mM IPTG; circles) conditions. Strains were sampled at 0, 45, 90, 135, 180 and 225 Minutes After Induction (MAI). Data from one out of three biological replicates are reported. C) Immunoblots of total protein extracts from *indrpsA* and *indPRPS1 E. coli* strains grown as described above sampled at 0, 45, 90, 135, 180 and 225 MAI, using S1 and PRPS1 specific antibodies. C.B.B.-stained gels are added as loading controls. Numbers show the relative protein abundance with respect to 0 MAI. Data from one out of three biological replicates are shown. Standard deviation was below 15%.

The possible functional homology between the two proteins was then investigated by introducing the *PRPS1* gene in *E. coli* cells, for over-expression and complementation assays. To experimentally test the ability of PRPS1 to complement S1 functions and to repress cell growth when over-accumulated in bacterial cells, both *rpsA* (encoding the S1 protein) and *PRPS1* coding sequences were cloned into pQE31-pREP4 plasmid system under the control of *pT5-lacO* promoter in pQE31. The plasmids were then introduced into the arabinose-dependent strain C-5699 (Briani *et al*., 2008), in which the chromosomal *rpsA* gene is transcribed from the *araBp* promoter, generating *indrpsA* and *indPRPS1* strains. Such systems provide a mechanism to deplete cells of the endogenous S1 protein in absence of arabinose and presence of glucose (Briani *et al*., 2008; Delvillani *et al*., 2011) and to modulate the expression of either *rpsA* or *PRPS1* genes cloned in pQE31 depending on IPTG concentration.

Both *indrpsA* and *indPRPS1* strains were cultured in three different conditions: Depletion (0.4% glucose and no IPTG), Complementation (0.4% glucose and 0.01 mM IPTG) and Overexpression (1% arabinose and 1 mM IPTG). The growth conditions for the complementation assay were experimentally optimized based on the growth of *indrpsA E. coli* strain. Cell growth was measured every 45 minutes up to 225 minutes after the induction (MAI, Fig. 5 B). At each time point cells were sampled, normalized on the optical density (OD_600_) value, and the total protein extract was used to detect the accumulation of S1 and PRPS1 proteins in both strains (Fig. 5 C). Under Depletion conditions, both strains showed almost completely impaired growth rate, due to the limited accumulation/absence of S1 protein. When grown in Complementation conditions, *indrpsA* cells were able to actively replicate and showed comparable amounts of S1 protein at each time point, indicating that the experimental conditions were properly set up to induce a wild-type-like S1 protein accumulation. On the other hand, *indPRPS1* cells failed to sustain growth in such conditions despite the gradual accumulation of PRPS1 protein, proving the inability of PRPS1 to complement the S1 function in *E. coli*. As expected, under Overexpression conditions, *indrpsA* strain growth was inhibited due to the excessive amount of S1 protein (Fig 5 B, C). Strikingly, the over-accumulation of PRPS1 protein was effectively repressing the bacterial growth and led to S1 protein depletion, too (Fig. 5 B, C). It is worth noting, that the overaccumulation of PRPS1 Arabidopsis protein was able to repress the growth of *E. coli* cells, comparably to S1 overaccumulation, even when the overexpression of either *rpsA* or *PRPS1* genes was achieved in the C-1a *E. coli* strain, devoid of the conditional depletion system of the endogenous S1 protein (Fig. S4 A). To better understand whether PRPS1 protein is capable of interacting with *E. coli* ribosomes under overexpression conditions, we sampled both *indrpsA* and *indPRPS1* cells at 90 MAI. Cell lysates were then fractionated into ribosome- unbound (supernatant, SN) and ribosome-bound (pellet, P) fractions and analysed via immunoblotting to detect either S1 or PRPS1 protein localisation. Interestingly, PRPS1 was retrieved in both ribosome-bound and -unbound fractions, similarly to S1 from *E. coli*, suggesting that the Arabidopsis PRPS1 can compete for the ribosome core with *E. coli* S1 protein (Fig. S4 B). Taken together, these data indicate that PRPS1 protein is able to inhibit *E. coli* growth when over-expressed in addition to the endogenous S1, whereas it is unable to functionally replace the *E. coli* endogenous S1 protein.

### PRPS1 accumulation is negatively regulated by chloroplast CLP protease complex

The phenotypes observed in Arabidopsis plants upon *PRPS1* overexpression indicate that the abundance of PRPS1 protein must be kept under a strict post-transcriptional control to prevent inhibition of protein synthesis and chloroplast damage (see Fig. 2-4). This notion is supported further by the inhibitory role of PRPS1 protein over-accumulation on *E. coli* cell growth (Fig. 5 and Fig. S4). In order to investigate the molecular mechanism responsible for controlling PRPS1 protein abundance, leaf discs were infiltrated with DEX supplemented with lincomycin (LIN), a specific inhibitor of plastid 70S ribosomes. Strikingly, a large accumulation of PRPS1 protein, most probably as result of PRPS1 degradation suppression, could be observed even after 24 hours from DEX infiltration, unlike the control sample (Fig. 6 A). Intriguingly, the most relevant chloroplast stromal protease is represented by the CLP complex, which is composed by several nuclear-encoded subunits and one plastid-encoded component, ClpP1, that is part of central proteolytic core (van Wijk, 2015; Llamas *et al*., 2017). To investigate whether the CLP complex could indeed be responsible for maintaining PRPS1 protein below levels that would otherwise cause damages to the chloroplast, we crossed *prps1-1* with mutants altered either in chloroplast protein translation, *prpl11-1*, or lacking two plastid chaperones required to feed CLP protease with protein substrates, *clpc1-1* and *clpd-1* (Pulido et al., 2016; Fig. 6 B). *prps1-1 prpl11-1* double mutant showed reduced growth rate (Fig. S5) and a slight decrease in photosynthetic efficiency (*F_V_/F_M_*) with respect to *prpl11-1* parental line. Similar genetic interactions were observed in *prps1-1 clpc1-1* double mutant, which showed a severe reduction in growth rate with respect to *prps1-1* and *clpc1-1* single mutants. Interestingly, *prps1-1 clpd-1* double mutant showed partially restored growth rate, close to wild-type-like levels, and a slight recovery of *F_V_/F_M_*parameter. Strikingly, the accumulation of PRPS1 protein increased about two- fold in all the double mutants tested (Fig. 6 C) with respect to *prps1-1*, further supporting the notion that plastid translation and the CLP protease complex play an important role in controlling PRPS1 abundance in the stroma of chloroplasts.

**Figure 6.**
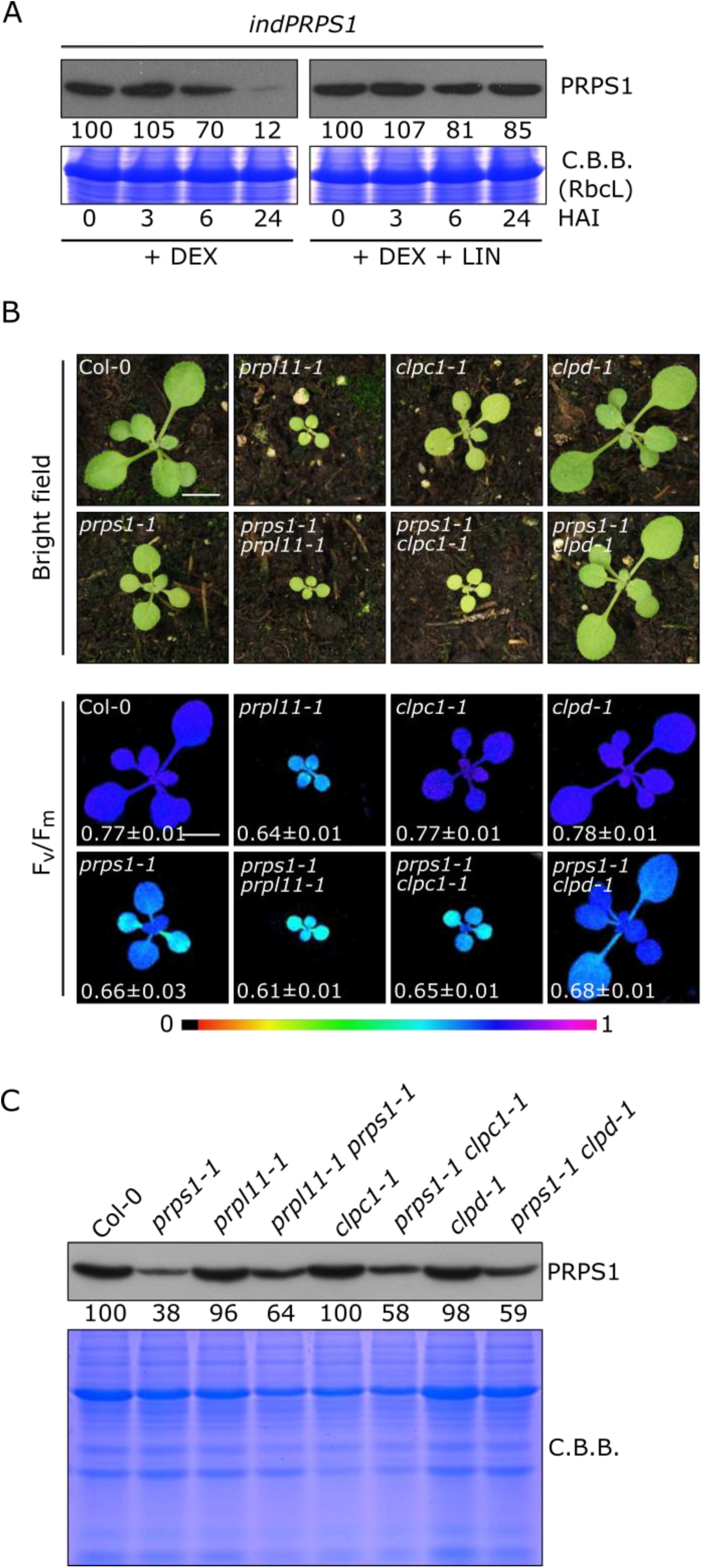
PRPS1 protein accumulation and the stromal CLP protease. A) Immunoblots of total protein extracts from *indPRPS1* leaf discs sampled at 0, 3, 6 and 24 HAI with 2 µM dexamethasone alone (+ DEX) or supplemented with 550 µM lincomycin (+ DEX + LIN). Filters were incubated with antibodies raised against PRPS1 protein. C.B.B.-stained gels are shown as loading control. Numbers below the immunoblots show the relative protein abundance with respect to 0 HAI (indicated as 100). Data are derived from one out of three biological replicates. Standard deviation was below 15%. B) Visible phenotypes and photosynthetic efficiency (*F_V_/F_M_*) of 16 Days After Sowing (DAS) seedlings grown on soil. On top (bright field) panel, visible phenotypes of Col-0 and *prps1-1*, *prpl11-1*, *clpc1-1*, *clpd-1*, *prps1-1 prpl11-1*, *prps1-1 clpc1-1* and *prps1-1 clpd-1* mutants are shown. On the bottom panel (*F_V_/F_M_*), the photosynthetic efficiency of each genotype is shown both as false colour imaging and average values ± standard deviations. Scale bar = 1 cm C) Immunoblot of total protein extracts from Col-0 and *prps1-1*, *prpl11-1*, *clpc1-1*, *clpd-1*, *prps1-1 prpl11-1*, *prps1- 1 clpc1-1* and *prps1-1 clpd-1* mutants obtained by using a PRPS1 specific antibody. C.B.B.-stained gels are shown as loading control. Numbers below the immunoblot show the relative protein abundance with respect to Col-0. Standard deviation was below 15%.

### Short-term induction of *PRPS1* and *PRPL4* gene expression induces different nuclear gene expression responses

The opposite behaviour of *indPRPS1* and *indPRPL4* plants in terms of protein pattern accumulation upon DEX-mediated induction of gene expression make them the ideal genetic material to investigate the primary nuclear gene expression response to changes in plastid ribosomal protein content. To this aim, a transcriptome analysis was performed on leaf discs harvested from *indPRPS1* and *indPRPL4* and vacuum infiltrated in either the absence or presence of DEX. In particular, total leaf RNA was extracted after 6 hours from infiltration, i.e., at the stage of maximal accumulation of *PRPS1* and *PRPL4* transcripts (see Fig. 4 A) and at the beginning of evident changes in protein pattern accumulation (see Fig. 4 B and C), and subjected to Illumina sequencing. Principal component analyses (PCA) of *indPRPS1* and *indPRPL4* samples showed a clear separation between the untreated and treated samples (Fig. S6), despite the short induction time. The EdgeR package was used to identify differentially expressed genes (DEGs, listed in Table S2), obtained by comparing data from *indPRPS1*-DEX with *indPRPS1+*DEX, and *indPRPL4-*DEX with *indPRPL4+*DEX (Fig. 7 A). Overall, 431 DEGs in *indPRPS1* upon DEX induction and 328 in *indPRPL4* were identified. Among them, 124 DEGs were in common between the two datasets. Next, the obtained DEGs were divided into up and down regulated genes both in *indPRPS1* and *indPRPL4* datasets, obtaining *indPRPS1*UP, *indPRPL4*UP, i*ndPRPS1*DOWN and *indPRPL4*DOWN lists, which were further compared to isolate unique DEGs in each group (Fig. 7 B, see also Table S2). As a result, among the up-regulated DEGs, 161 were uniquely found in *indPRPS1* (Table S2 A) and 159 DEGs in *indPRPL4* (Table S2 B), while 107 DEGs were up-regulated in both lines (Table S2 C). Among the down-regulated DEGs, 146 were unique for *indPRPS1* line (Table S2 D), 45 unique DEGs for *indPRPL4* (Table S2 E) and only 14 were commonly down-regulated (Table S2 F). Furthermore, among the 149 down-regulated DEGs in *indPRPS1*, 3 were upregulated in *indPRPL4* (Table S2 G).

**Figure 7.**
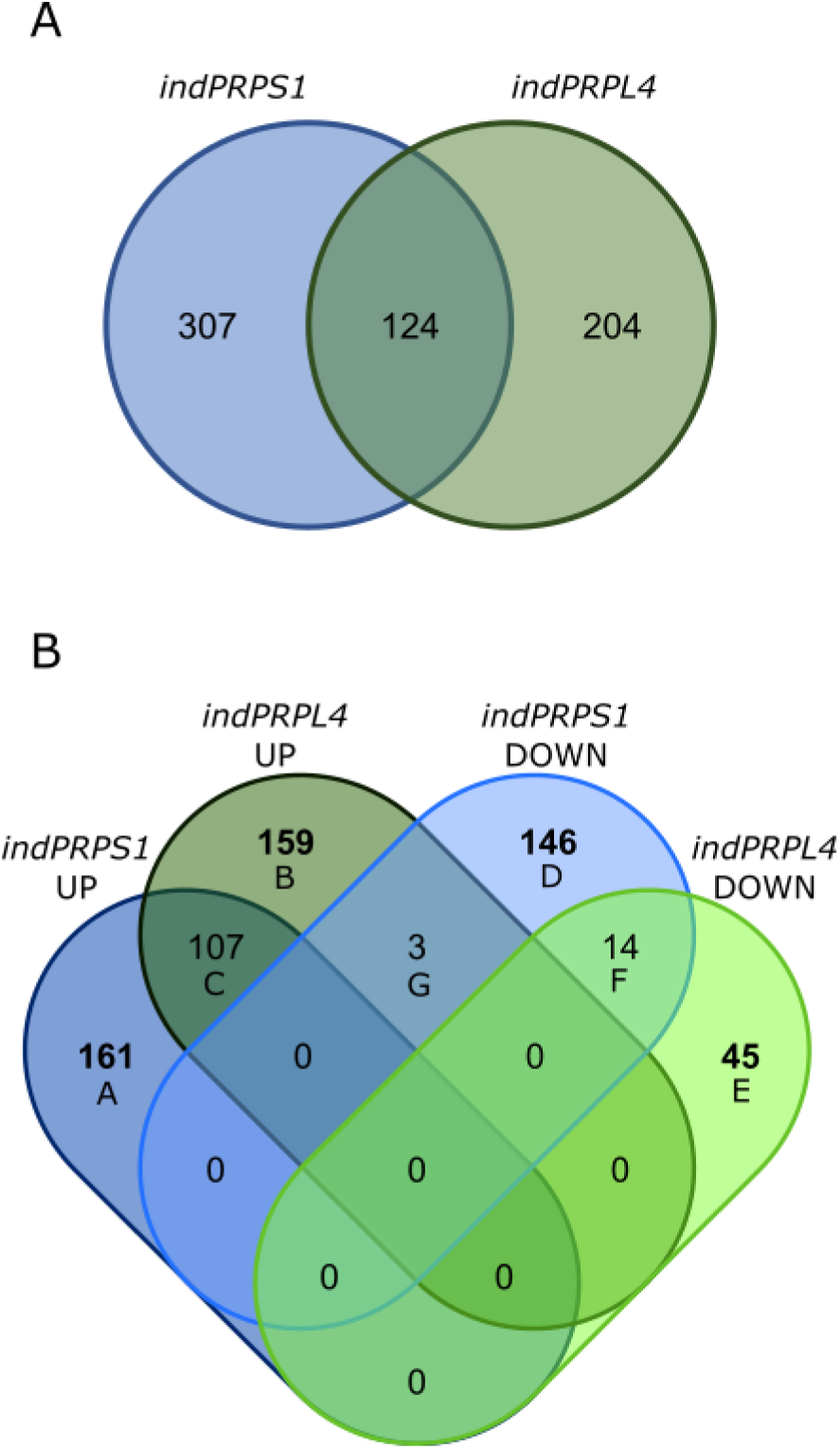
Comparison of RNAseq transcriptomes from *indPRPS1* and *indPRPL4* lines infiltrated or not with DEX. A) Venn diagram showing the number of DEGs found in *indPRPS1* and *indPRPL4* upon induction. B) Venn diagram showing the distribution of DEGs resulted from the comparison of *indPRPS1*UP, *indPRPL4*UP, *indPRPS1*DOWN and *indPRPL4*DOWN lists. Unique DEGs in each group are in bold. Group A: Unique up-regulated DEGs found in *indPRPS1*; Group B: Unique up-regulated DEGs found in *indPRPL4*; Group C: Common up-regulated DEGs found in *indPRPS1* and *indPRPL4*; Group D: Unique down-regulated DEGs found in *indPRPS1*; Group E: Unique down- regulated DEGs found in *indPRPL4*; Group F: Common down-regulated DEGs found in *indPRPS1* and *indPRPL4*; Group G: up-regulated DEGs found in *indPRPL4* but down-regulated in *indPRPS1*.

To investigate the biological functions activated or repressed by the overexpression of either *PRPS1* or *PRPL4* genes, the unique DEGs found up- or down-regulated in *indPRPS1* or *indPRPL4* were analysed with agriGO v2.0 online tool (Tian *et al*., 2017). Next further analysis by REVIGO (Supek et al., 2011) was preformed to retrieve Biological Process Gene Ontology terms and group them into functional categories (Fig. 8; Table S3, S4, S5). The 161 DEGs up-regulated in *indPRPS1* produced strongly enriched GO terms associated with responses to endogenous factors or involvement in flowering and seeds production, such as “photoperiodism, flowering” (GO:0048573), “response to karrikin” (GO:0080167), “vegetative to reproductive phase transition of meristem” (GO:0010228) or “cellular response to hormone stimulus” (GO:0032870) (Fig. 8 A; see also Table S3). On the other hand, the repressed biological functions found in *indPRPS1* were related to the production of secondary metabolites, defence against herbivores and oxidative stress such as “glucosinolate biosynthetic process” (GO:0019761), “sulfur compound biosynthetic process” (GO:0044272), “cell redox homeostasis” (GO:0045454) and “response to wounding” (GO:0009611), to cite a few of them (Fig. 8 B; see also Table S4). Interestingly, the high enrichment in GO terms associated with flowering and reproductive phase transition is in good agreement with the anticipated flowering phenotype, calculated as number of rosette leaves at bolting, observed when *PRPS1* transcripts are overexpressed (Fig. 9). In particular, in the case of plants cultivated on soil under growth chamber conditions, Arabidopsis Col-0 bolted with an average of about 10-11 rosette leaves while the *oePRPS1* line showed an anticipated flowering time, bolting at the stage of 8 rosette leaves, comparable to *prps1-1* behaviour (Fig 9 A, B). The same anticipated flowering time occurred under short day growth conditions, with the wild-type bolting at 54 leaves while the overexpressing line flowered at 37 rosette leaves on average (Fig 9 C). A similar observation was made when *indPRPS1* line was grown on MS medium supplemented with 1% (w/v) sucrose and DEX, in which its number of leaves at bolting was 5, similarly to *oePRPS1* (Fig. 9 D). Instead, when *indPRPS1* was grown on MS medium without DEX, the number of leaves at bolting was about 6-7, comparable to what observed in *indPRPL4* and Col-0 plants.

**Figure 8.**
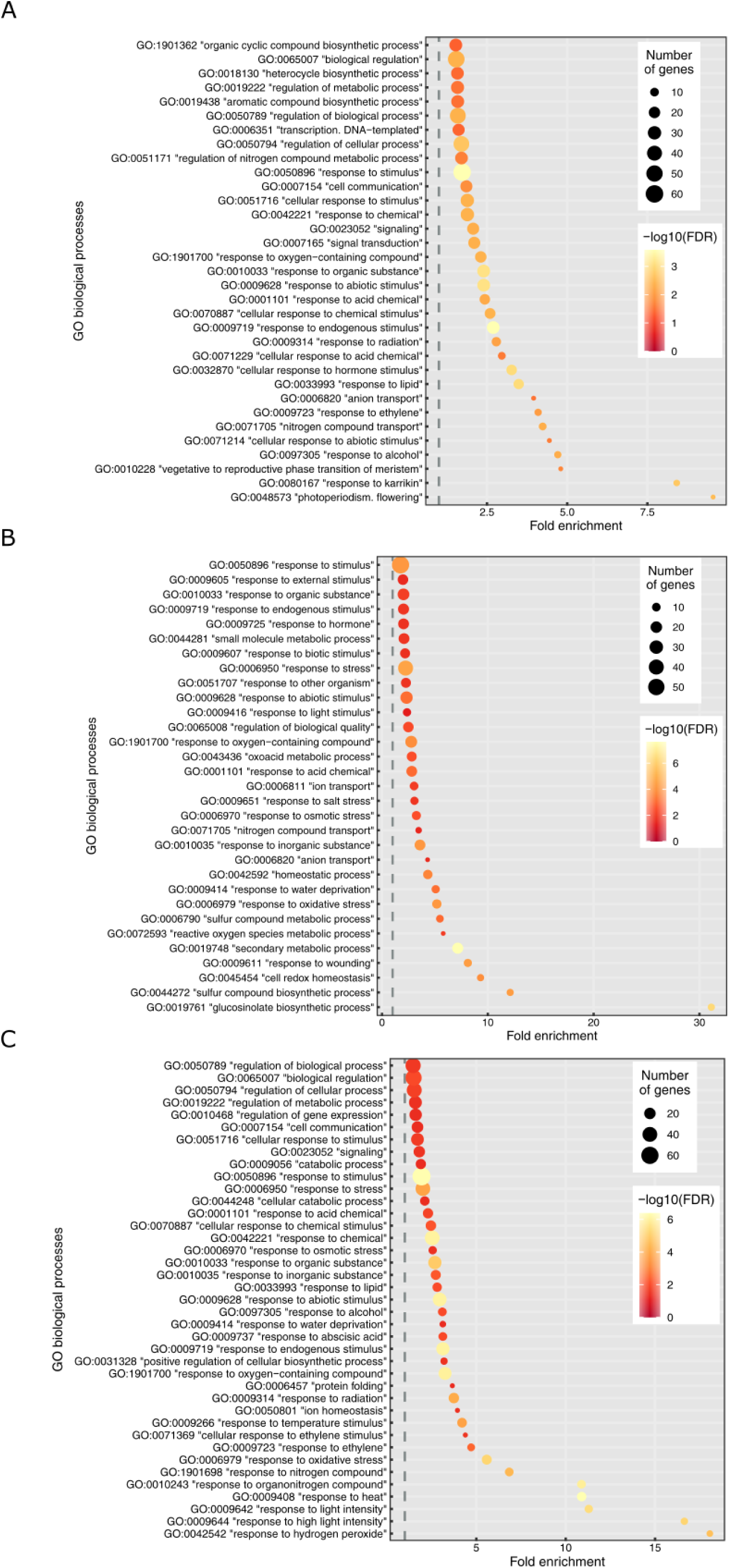
Analysis of the biological functions activated or repressed by the short-term overexpression of *PRPS1* and *PRPL4* genes. Significantly enriched GO terms identified through agriGO v2.0 and REVIGO online tools retrieved from A) unique up-regulated DEGs found in *indPRPS1*; B) unique down-regulated DEGs found in *indPRPS1;* C) unique up-regulated DEGs found in *indPRPL4*.

**Figure 9.**
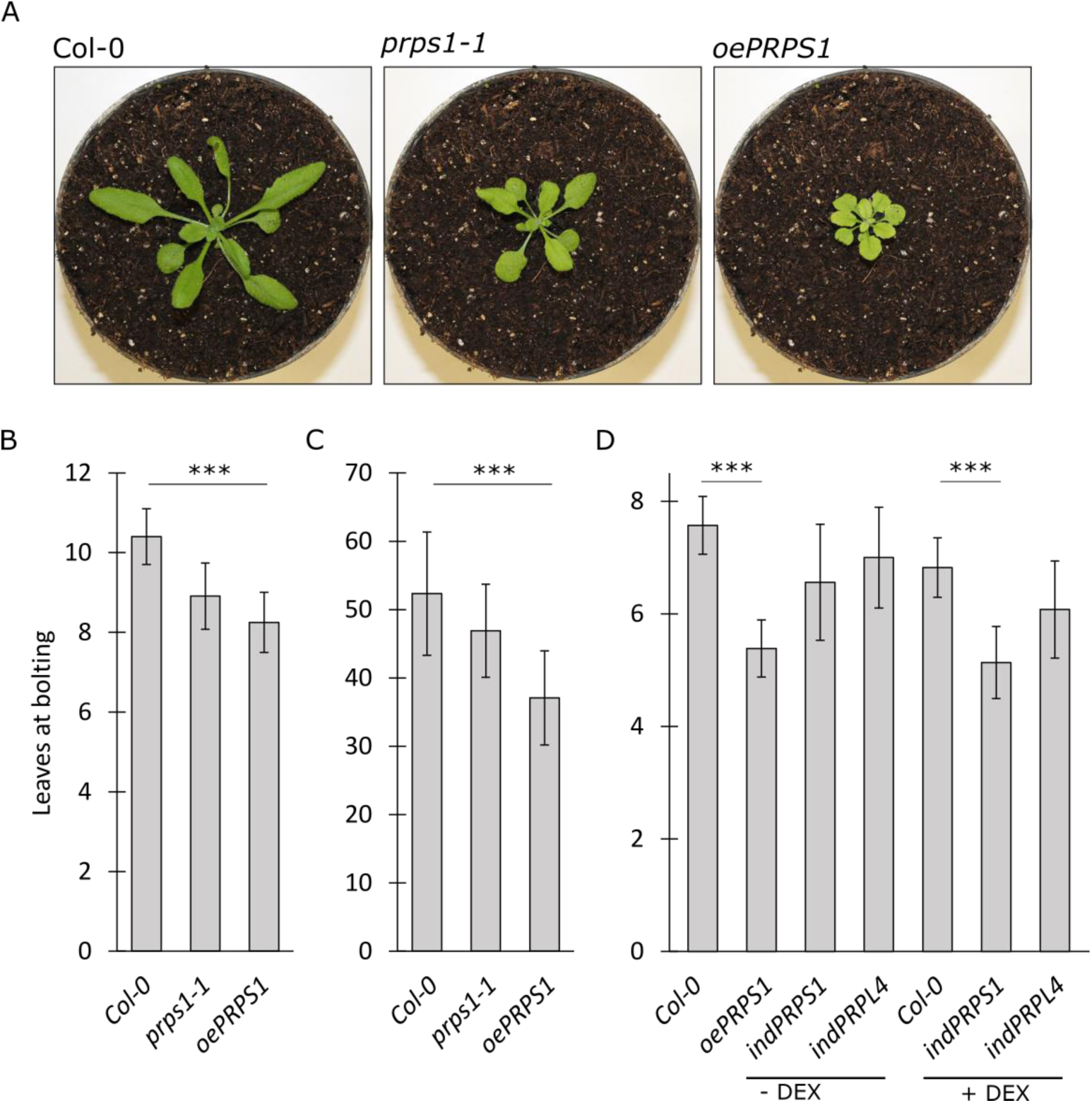
Flowering time determination in Col-0, *prps1-1*, *oePRPS1*, *indPRPS1* (± DEX) and *indPRPL4* (± DEX) lines. A) Pictures of representative plants at bolting grown on soil in long day conditions. B) Average numbers ± standard deviations of leaves at bolting observed in Col-0, *prps1- 1* and *oePRPS1* plants grown on soil in long day conditions. C) Average numbers ± standard deviations of leaves at bolting observed in Col-0, *prps1-1* and *oePRPS1* plants grown on soil in short day conditions. D) Average numbers ± standard deviations of leaves at bolting observed in Col-0, *oePRSP1*, *indPRPS1* and *indPRPL4* plants grown on MS medium devoid (- DEX) or supplemented (+ DEX) with 2 µM dexamethasone in long day conditions. Statistical significance was calculated via Student’s t-test (*** indicates *P* < 0.001).

Conversely, the analysis of the activated biological processes in *indPRPL4* showed a strong enrichment in GO terms associated with abiotic stress responses and protein homeostasis such as “response to hydrogen peroxide” (GO:0042542), “response to high light intensity” (GO:0009644), “response to heat” (GO:0009408), “response to oxidative stress” (GO:0006979) and “protein folding” (GO:0006457) (Fig. 8 C; see also Table S5). These activated cellular pathways are in agreement with the observed increased levels of CLPB3, HSP90-1 and HSC70-4 proteins in *indPRPL4* line upon induction with DEX (see Fig. S7). In particular, CLPB3 protein abundance and even more the abundance of HSP90-1 and HSC70-4 proteins largely increased already at 3 HAI in *indPRPL4* leaf disks, unlike in *indPRPS1* samples, possibly generated by the overload of the plastid folding machinery caused by PRPL4 abundance and the consequent activation of chloroplast UPR. Interestingly, the unique down-regulated DEGs found in *indPRPL4* leaf disks resulted in no significantly enriched GO terms.

### Short-term induction of *PRPS1* and *PRPL4* gene expression induces different changes in proteomic profiles

To further investigate the molecular responses following the DEX-mediated induction of *PRPS1* and *PRPL4* genes, a proteomic analysis was performed on leaf tissue harvested from *indPRPS1* and *indPRPL4* plants and vacuum infiltrated either in the absence or presence of DEX and sampled, as in the case of transcriptome analysis, after 6 hours of induction. PCA analysis revealed that dexamethasone-mediated induction led to significant changes also at proteomic level (Fig. S8). Comparing induced (+DEX) and control groups (-DEX), the abundances of 58 and 77 proteins were altered in *indPRPS1* and *indPRPL4* samples, respectively (T Student’s test, FDR < 0.05; listed in Table S6). The minimal overlapping between the two datasets (only two common proteins) confirms the different cellular responses following the induction of *PRPS1* and *PRPL4* genes (Fig. 10 A). In particular, 27 and 31 proteins were up- and down-accumulated in *indPRPS1* leaf discs, whereas 45 and 32 proteins were up- and down-accumulated in *indPRPL4* samples, respectively (Fig. 10 B).

**Figure 10.**
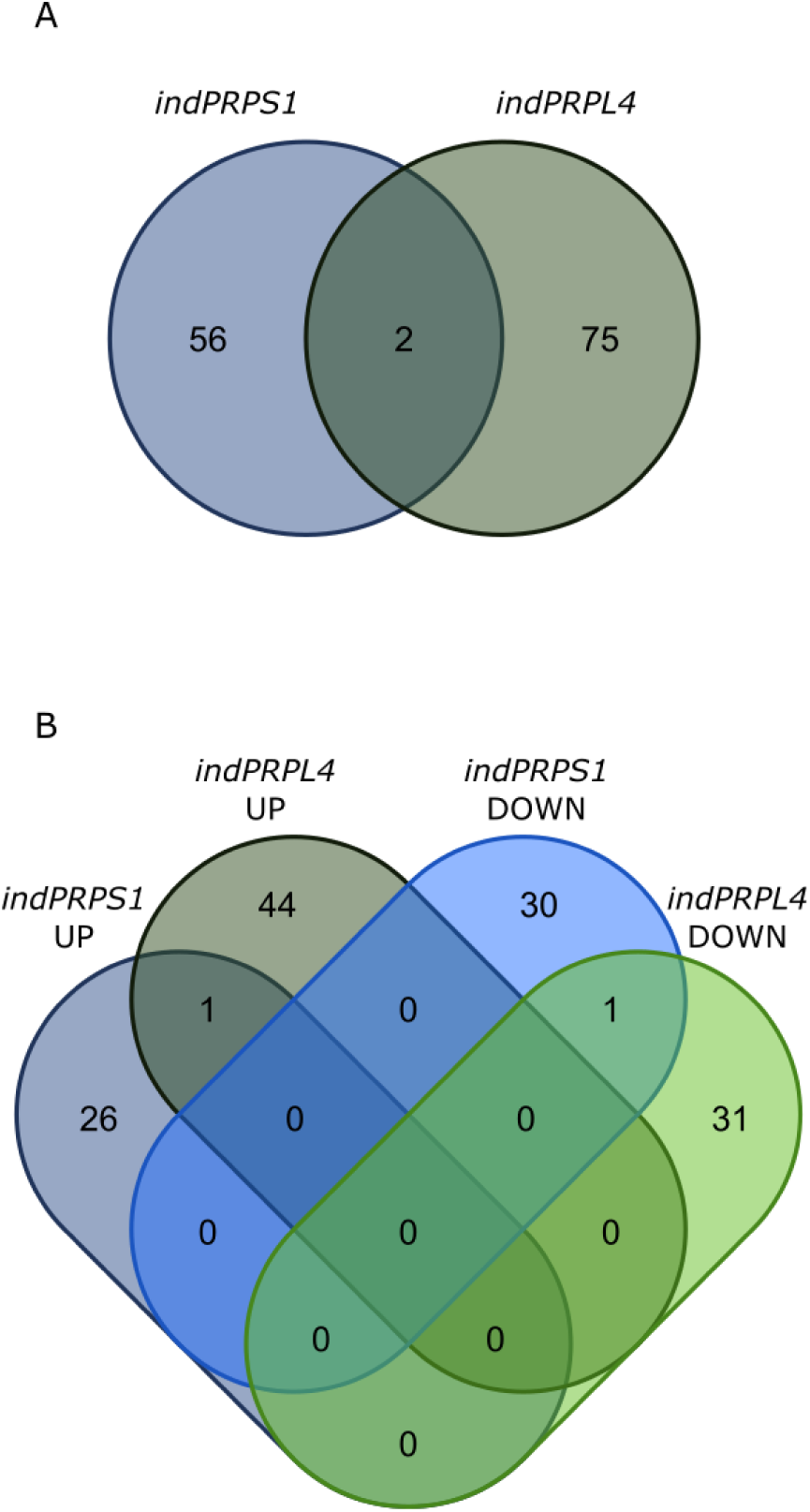
Comparison of proteome from *indPRPS1* and *indPRPL4* lines infiltrated or not with DEX. A) Venn diagram showing the number of DAPs found in *indPRPS1* and *indPRPL4* upon induction. B) Venn diagram showing the distribution of DAPs resulted from the comparison of *indPRPS1*UP, *indPRPL4*UP, *indPRPS1*DOWN and *indPRPL4*DOWN lists.

GO enrichment analysis of differentially abundant proteins (DAPs) found in *indPRPS1* samples revealed over-represented GO terms only among the down-accumulated proteins, probably due to the small size of the data set. The enriched Biological Process terms were “translational elongation” (GO:0009658) and “chloroplast organization” (GO:0006414) (Table S7). Among the enriched Cellular Component terms, “ribosome-associated quality control (RQC) complex” (GO:1990112), “transcriptionally active chromatin” (GO:0035327) and “chloroplast stroma” (GO:0009570) were found (Table S8). All DAPs were grouped based on their GO annotations (Fig. 11). Proteins up-regulated in response to *PRPS1* induction were mainly involved in “anatomical structure development” (GO:0048856), “catabolic process” (GO:0009056) and “response to light stimulus” (GO:0009416) (Fig. 11 A). In addition, the up-accumulated proteins in *indPRPS1* plantlets were mainly located in “nucleus” (GO:0005634) and “cytoplasm” (GO:0005737) (Fig. 11 B). Down- regulated DAPs produced GO annotations such as “biosynthetic processes” (GO:0009058), “RNA binding” (GO:0003729), “translation” (GO:0006412) and “post-embryonic development” (GO:0009791) (Fig. 11 A). Among down-regulated proteins detected in *indPRPS1* samples we found the regulator of fatty-acid composition 3 (RFC3; AT3G17170) which plays an important role in the plastid rRNA processing (Nagashima *et al*., 2020). Many of these down-regulated proteins were located in chloroplast and nucleus (Fig. 11 B).

**Figure 11.**
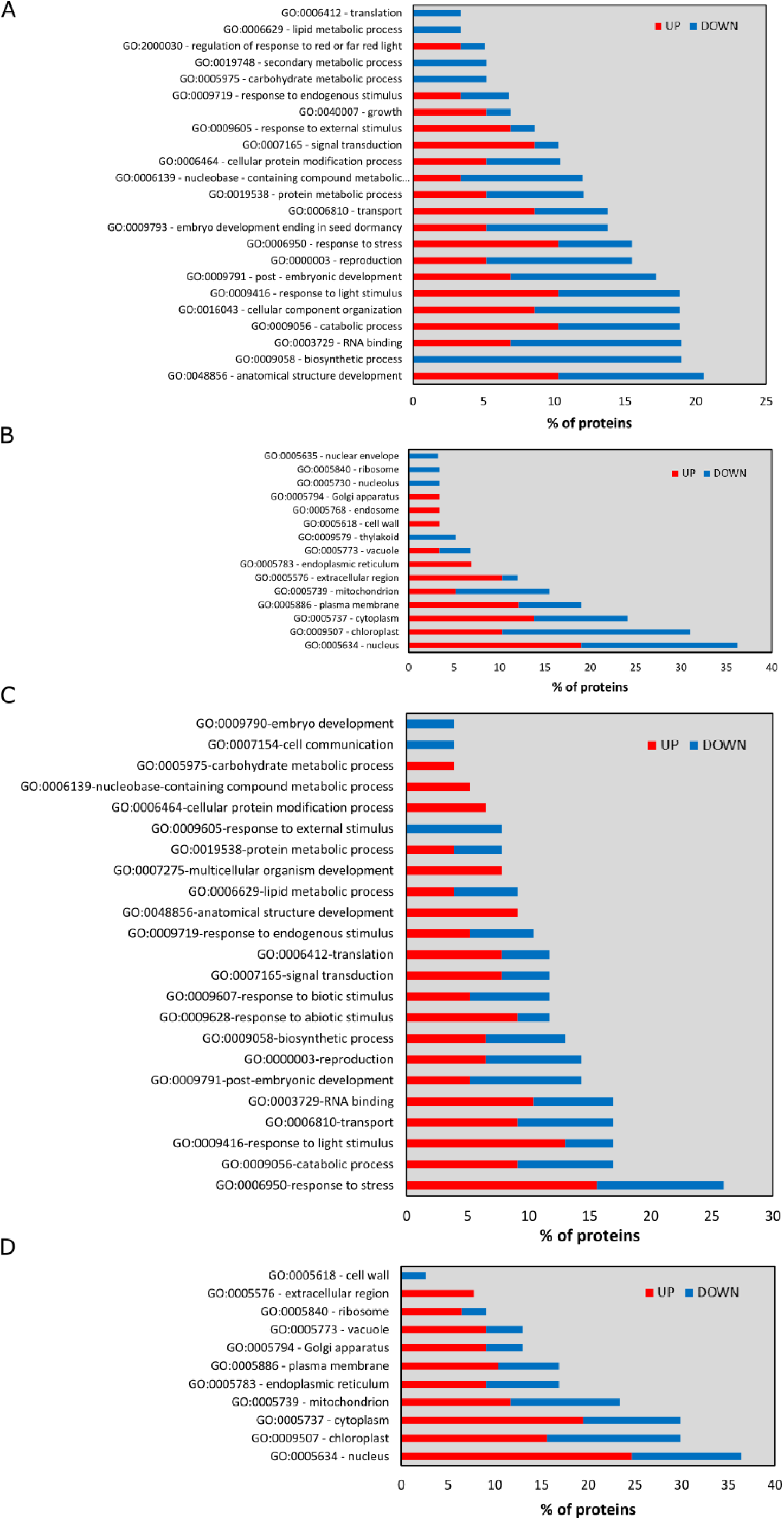
Protein categorization based on Biological Process GO terms of DAPs found in *indPRPS1* (A) and *indPRPL4* (C) samples and based on Cellular Content GO terms found of *indPRPS1* (B) and *indPRPL4* (D) samples. Blue and red bars correspond to proteins that were up or down accumulated, respectively.

The significantly enriched GO terms retrieved by analysing the up-regulated proteins detected in *indPRPL4* samples were mainly associated with mitochondrial electron transport chain and mitochondrial ribosomes (Table S9). Moreover, GO annotation revealed that several DAPs in *indPRPL4* seedlings belong to “response to stress” (GO:0006950), as also detected by the transcriptome analysis (Fig. 11 C). The GO annotation tool allowed to locate DAPs in *indPRPL4* plants especially in the nucleus, chloroplast and cytoplasm (Fig. 11 D). Overall, our results, in agreement with transcriptomic analysis, point out very specific proteomic changes following the alteration of the two ribosomal proteins PRPS1 and PRPL4. In fact, the induction of *PRPS1* leads to a general repression of translation, transcription and chloroplast organization. Conversely, the increased amount of PRPL4 protein promotes the cellular stress response, leading to a positive regulation of proteins involved in RNA metabolism and translation, with a major contribution also given by components of the mitochondrial metabolism.

## Discussion

### Altered *PRPS1* expression impairs chloroplast biogenesis

*PRPS1* is a nuclear gene encoding the S1 protein of the plastid 30S small ribosomal subunits. Reduced expression of *PRPS1* gene results in diminished plastid translation and decreased photosynthetic efficiency (Romani *et al*., 2012; Yu *et al*., 2012), while its disruption arrests embryo development at the globular stage in Arabidopsis (see Fig. 1) and prevents greening of seedlings in rice (*albino seedling lethality 4*; Zhou et al., 2021). Clearly, these observations indicate that *PRPS1* is an essential gene required during early stages of chloroplast biogenesis, as reported in the case of other essential nuclear genes coding for plastid ribosomal proteins (Bryant et al., 2011; Lloyd and Meinke, 2012; Romani et al., 2012). The apparent discrepancy between the early block of embryo development in Arabidopsis and the arrest of seedling greening in rice is in agreement with previous findings (Hess *et al*., 1994; Zubko & Day, 1998; Ostheimer *et al*., 2003; Asakura & Barkan, 2006) and compatible with the essential nature of fatty acid biosynthesis during chloroplast biogenesis. In particular, the lack of the plastid-encoded accD-subunit of the multimeric acetyl-CoA carboxylase required for fatty acid biosynthesis is responsible for the lethality of Arabidopsis embryos defective in plastid translation (Bryant *et al*., 2011). In contrast, grass species contain a plastid-located monomeric acetyl-CoA carboxylase that, differently from Arabidopsis, is encoded in the nucleus and translated in the cytosol (Schulte *et al*., 1997; Chalupska *et al*., 2008). Therefore, fatty acid biosynthesis (and embryogenesis) can continue even when plastid protein synthesis is affected in these species.

Intriguingly, the reduced accumulation of S1 protein can also be obtained when *PRPS1* gene expression, and the consequent transcript accumulation, is both constitutively increased in *oePRPS1* and *indPRPS1* + DEX seedlings (Fig. 2) and induced in *indPRPS1* leaf discs for 24 hours (Fig. 4). This finding, together with the very limited increase in transcript accumulation observed in *oePRPS1* and *indPRPS1* + DEX seedlings, indicates the existence of post-transcriptional regulatory mechanisms aimed to prevent PRPS1 over-accumulation, as shown previously (Yu *et al*., 2012). The strict control on *PRPS1* gene expression and protein accumulation appears to be rather specific and certainly it does not apply to *PRPL4*, a nuclear gene encoding a core subunit of the 50S plastid ribosome, also essential for embryo development and plant viability (Bryant *et al*., 2011; Romani *et al*., 2012). In fact, *oePRPL4* and *indPRPL4* + DEX lines were able to accumulate 25-30 times more transcripts, together with the double amount of PRPL4 protein, in comparison to Col-0 and *indPRPL4* - DEX controls (Fig. 2, Fig. 4).

Furthermore, the constitutive over-expression of *PRPS1* gene, driven in 16 DAS *oePRPS1* and *indPRPS1* + DEX plantlets, resulted in the impairment of chloroplast differentiation and physiology, leading to a virescent phenotype visible in the emerging leaves and in the younger portions of the leaves, corresponding to the tissue proximal to the petiole. In particular, transmission electron microscopy (TEM) observations of the chlorotic tissues revealed the presence of cells displaying misshapen chloroplasts with altered thylakoid membrane ultrastructure and large plastoglobuli in the stroma (Fig. 3), reported to be associated with the degradation of chlorophylls and thylakoids in response to abiotic and biotic stresses or during senescence (Rottet *et al*., 2015; Van Wijk & Kessler, 2017; Zechmann, 2019). Moreover, the presence of vesicles budding from chloroplasts and containing electron dense material, together with the observation of entire round chloroplasts inside the vacuole (Fig. 3), indicates the ongoing chloroplast degradation, similarly to what reported in literature (Woodson, 2016; Zhuang and Jiang, 2019; Tadini et al., 2020; Jeran et al., 2021). Coherently, the relative expression of *ATI1* and *ATG8f* genes, involved in ATG-dependent micro-autophagy, was enhanced upon PRPS1 attempted over-accumulation (Fig. 3, Fig. S1 A), whereas the expression of genes involved in ATG-independent micro-autophagy was mildly stimulated in *oePRPS1* (Fig. S1 B) (Lemke *et al*., 2021). Overall, these observations indicate that the reduced accumulation PRPS1 severely jeopardizes chloroplast integrity during leaf development, leading to the dismantling of damaged and misshapen chloroplasts with the final aim to remove reactive oxygen species–producing chloroplasts and redistributing nutrients to other tissues (Woodson, 2022).

### A fully functional proteostasis machinery is needed to control PRPS1 accumulation in chloroplast stroma

All our attempts to increase the abundance of PRPS1 protein failed (Fig. 2 and Fig. 4; Tadini *et al*., 2016) and, together with that, the accumulation of other plastid ribosomal proteins, such as PRPL4 and PRPS5, was reduced, as observed in *indPRPS1* + DEX leaf discs (Fig. 4), indicating that the decreased abundance of PRPS1 protein has a deleterious effect on plastid ribosome stability. In agreement with that, the pulse-labelling experiment conducted on *indPRPS1* leaf material infiltrated with DEX for 6 hours showed a severe inhibition of plastid protein translation and a larger fraction of PRPS1 protein freely associated to mRNA and not bound to actively translating ribosomes (Fig. 4). These findings explain the defects in leaf greening observed in the different Arabidopsis lines (Fig. 1 and Fig. 2) and support the role of PRPS1 as a stringently regulated translation factor rather than a “real” ribosomal protein, given its weakly and reversible association with the 30S subunit, similarly to previous observations in *E. coli* (Delvillani *et al*., 2011).

S1 protein is the closest PRPS1 homologue in *E. coli*, and it is encoded by the essential gene *rpsA* (Kitakawa & Isono, 1982). To gain possible insights on PRPS1 function, we attempted to rescue *E. coli* cell lethality due to S1 depletion and to phenocopy the bacteriostatic effects of S1 overaccumulation, by modulating the level of PRPS1 protein in *E. coli* cells (Fig. 5). Our data clearly showed that PRPS1 is not able to functionally replace the endogenous S1, as cells depleted of *rpsA* but with moderate amount of PRPS1, failed to grow (Fig. 5). According to previous studies, S1 protein exerts its functions in relationship with the specializations of its six S1 domains (Salah *et al*., 2009). The interactions with the ribosome relies on S1 domains 1 and 2 (Giorginis & Subramanian, 1980), whereas the ability to bind mRNAs has been associated with the S1 domains 3, 4 and 5 (Subramanian, 1983). As for domain 6, if removed together with S1 domain 5, the initiation of translation is hampered (Boni *et al*., 2000; Salah *et al*., 2009) and together with domain 4, it is implicated in ribosome dimerization and hibernation in stationary phase (Beckert *et al*., 2018). According to our *in silico* analysis, S1 domains 1 and 2 of PRPS1 appeared to be more similar to the respective domains 1 and 2 of S1, while domain 3 clustered together with domains 3, 4 and 5 of S1 (Fig. S3). The fact that PRPS1 is a smaller protein and possesses three out of the six S1 domains identified in S1, of which none could be associated with those required for translation initiation in *E. coli*, could explain the inability of PRPS1 to functionally replace S1 protein. Nonetheless, PRPS1 overaccumulation blocked cells growth as much as S1 (Fig. 5, Fig. S4). It has been shown that S1 over accumulation inhibits translation since the excess of “free” S1 interacts with mRNAs, preventing the ribosome loading (Delvillani et al., 2011; Boni et al., 2001, Skorski et al., 2006; Skouv et al., 1990). However, PRPS1 was found both in ribosome-bound and -unbound fractions, suggesting that, although PRPS1 can interact with the ribosome core in *E. coli* cells, this interaction is rather unfruitful. In this scenario PRPS1 would be able to inhibit *E. coli* growth by competing with the endogenous S1 protein and inhibiting ribosome activity (Fig. S4). Nevertheless, PRPS1 is likely capable of binding *E. coli* mRNAs with the S1 domains 2 and 3, since *E. coli* and plastid mRNAs share similar 5’ UTRs with AU-rich sequences (Hirose & Sugiura, 2004), making *E. coli* transcripts inaccessible to ribosomes and acting as a negative modulator of translation initiation. The inhibitory role of PRPS1 protein on *E. coli* protein synthesis is corroborated further by the fact that the accumulation of PRPS1 upon overexpression reaches very rapidly the plateau level and concomitantly leads to the decreased accumulation of S1 protein over time (Fig. 5 C), mimicking the role of S1 as feedback effector of its own regulation at the translational level (Skouv *et al*., 1990).

Such regulatory mechanism has a different spatial constraint in photosynthetic eukaryotes due to the physical separation of the nuclear/cytosolic compartments, where the *PRPS1* transcripts and the precursor protein are synthesized, and the chloroplast stroma where the mature PRPS1 protein plays its functions. While the limited accumulation of *PRPS1* transcripts observed upon constitutive expression allows us to hypothesize the existence of a cytosolic post-transcriptional regulatory mechanism (Wu *et al*., 2019), our data strongly support the activation of a chloroplast stroma post- translational regulatory mechanism, in response to *PRPS1* overexpression, mediated by the plastid proteostasis machinery (Fig. 6). In chloroplasts, the major soluble stromal protease is the CLP complex, composed of multiple nuclear-encoded subunits with the addition of ClpP1 subunit, the only one to be plastid-encoded (van Wijk, 2015). Consequently, ClpP1 abundance is susceptible to genetic defects in plastid gene expression or to drugs inhibiting plastid translation (Llamas *et al*., 2017). The co-infiltration of DEX with the plastid translational inhibitor lincomycin allowed a larger accumulation of PRPS1 protein (Fig. 6 A). Accordingly, an increased PRPS1 protein accumulation (Fig. 6 C) could be achieved by introgressing the *prps1-1* knock-down allele into *prpl11-1* genetic background, in which the chloroplast translation is reduced (Pesaresi *et al*., 2001). This is also in line with the restoration of PRPS1 protein accumulation observed in *prps1 gun1* and *prps1 rh50* double mutants, as GUN1 stimulates the activity of the Nuclear-Encoded Polymerase (NEP) that, among other plastid house-keeping genes, is responsible of the transcription of *clpP1*, while RH50 is involved in plastid ribosome assembly and plastid translation (Tadini *et al*., 2016, 2020c,a,b; Paieri *et al*., 2018). Intriguingly, GUN1 was also found to physically interact with the plastid protein homeostasis machinery, including ClpC subunits (Tadini *et al*., 2016). Furthermore, a comparable increase in PRPS1 levels were also observed in the double mutants *prps1-1 clpc1-1* and *prps1-1 clpd- 1* (Fig. 6 C), in which the two plastid chaperones required by CLP protease to interact with the substrates are missing (Pulido *et al*., 2016). These pieces of evidence strongly point towards CLP protease complex as one of the main regulators of PRPS1 protein abundance in Arabidopsis chloroplasts.

Our findings are also in agreement with previous reports. In particular, by combining transcriptomic and proteomic analyses, Wu et al., 2019 were able to show that plastid ribosomal proteins are regulated post-translationally, suggesting a protein degradation-based mechanism. Moreover, in *Chlamydomonas*, *CreS1* expression is induced by light, while CreS1 protein levels remain constant, pointing also in this case to the post-translational regulation of CreS1 abundance (Merendino *et al*., 2003).

On the other hand, the constitutive over-accumulation of PRPL4 protein upon induction didn’t affect neither the plastid translation nor the chloroplast ultrastructure (Fig. 2, Fig. 3, Fig. 4).

Interestingly, chaperones both resident in the cytoplasm and in plastids were progressively up- regulated as PRPL4 levels increased, indicating the activation of protein homeostasis mechanisms to cope with the increased protein amount (Fig. S7). These different responses to the overexpression of two plastid ribosomal proteins could be due to their intrinsic features and activities, having PRPS1 mRNA-binding properties which, if not kept under strict control, would have dramatic consequences on chloroplast functionality.

### *PRPS1* overexpression promotes early flowering while PRPL4 overaccumulation triggers cpUPR

The switch from vegetative growth to reproductive growth is a pivotal event in plant development, mostly dependent on the environmental stimuli such as day length (photoperiodic flowering) and temperature (vernalization) that plants perceive to determine the proper timing to assure successful reproduction, and ultimately the survival of the species (Freytes *et al*., 2021). However, plants subjected to a variety of stressful conditions can anticipate their flowering through a new category of flowering response, known as stress-induced flowering, aimed to guarantee species survival when they cannot adapt to unfavourable conditions (Takeno, 2016). Although the mechanistic details behind the stress-induced flowering are still not fully understood, hormones seem to be at least partially involved in these pathways as they are produced under stress and regulate gene expression to cope with it (Takeno, 2016; Kazan & Lyons, 2016). This seems to be the case of plants subjected to prolonged *PRPS1* overexpression and characterised by an early flowering phenotype regardless of day length (Fig. 9). In these plants, the stress seems to be caused by the reduced chloroplast protein synthesis and the consequent accumulation of damaged proteins and, possibly, by low sucrose concentration due to the reduced photosynthetic performance (see Fig. 2; Bolouri Moghaddam and Van den Ende, 2013). Consistently, the transcriptomic profile of Arabidopsis lines characterised by the short-term induction of *PRPS1* gene highlighted that the early nuclear gene expression response to PRPS1 imbalance in chloroplasts is characterised by a robust enrichment of up-regulated genes associated with the reproductive phase transition and the cellular response to hormones, such as “photoperiodism, flowering”, “vegetative to reproductive phase transition of meristem” and “cellular response to hormone stimulus” (Fig. 8 and Table S3), and by the concomitant repression, amongst others, of genes involved in “cell redox homeostasis”, “response to wounding”, “response to oxidative stress”, “response to water deprivation”. Accordingly, the circadian clock regulators *RVE8* and *CO*, the *IDD8* transcription factor and the Gibberellic acid enzyme *ga3ox1*, all positive modulators of flowering, are among the up-regulated genes. On the other hand, the tetratricopeptide thioredoxin-like protein *TTL4*, required for osmotic stress tolerance, *BASS5* and *BCAT4*, involved in glucosinolate biosynthesis, *PTR3*, required for defences against pathogens, and the transcription factor *NAC019*, involved in the response to dehydration, were found to be down- regulated upon *PRPS1* induction (Tran *et al*., 2004; Schuster *et al*., 2006; Karim *et al*., 2007; Sawada *et al*., 2009; Lakhssassi *et al*., 2012). Interestingly, the mass spectrometry analysis of the tissue overexpressing *PRPS1* detected the increased accumulation of far-red insensitive 219 (FIN219; AT2G46370) and the phytochrome associated protein phosphatase 2C (PAPP2C; AT1G22280), both directly involved in the response to red, or far red light and in flower development (Hsieh *et al*., 2000; Phee *et al*., 2008); while chloroplast-located proteins involved in “RNA metabolism” and “translation” where down-regulated.

In contrast, the results obtained from the transcriptomic and proteomic analyses performed on *indPRPL4* + DEX leaf discs showed markedly different results (Fig. 7-8, Fig. 10-11). The overaccumulation of PRPL4 promoted abiotic stress responses and protein homeostasis such as “response to hydrogen peroxide”, “response to high light intensity”, “response to heat”, “response to oxidative stress” and “protein folding” (Fig. 8 C). In particular, transcription factors involved in stress responses such as *DREB2A*, *DREB2C* and *WRKY26* (Sakuma *et al*., 2006; Chen *et al*., 2010; Li *et al*., 2011), were found to be upregulated, together with proteins directly involved in proteostasis maintenance, such as small HSPs, CLPB1 cytosolic unfoldase and the co-chaperone HOP3 (Sun *et al*., 2001; Mishra & Grover, 2016; Fernández-Bautista *et al*., 2017; Toribio *et al*., 2020). Interestingly, *GOLS1* gene coding for a key enzyme of raffinose family sugar synthesis, shown to enhance plant resistance to oxidative damage, was also up-regulated (Panikulangara *et al*., 2004; Nishizawa *et al*., 2008). These findings together with the increased accumulation of CLPB3, HSP90-1 and HSC70-4 proteins over time (see Fig. S7), corroborate the hypothesis that PRPL4 over-accumulation contributes to generate pressure on the plastid folding machinery that, in turns, activates retrograde signalling pathways aimed at triggering a chloroplast-related UPR. It can be envisaged that the activated cpUPR contributes to the limited increase of PRPL4 protein abundance in Arabidopsis chloroplasts, i.e., two folds higher upon 24 HAI with respect to 0 HAI, despite the 30-fold increase of PRPL4 transcripts (Fig. 4 A and C).

Taken together these observations highlight the activation of different nuclear and cellular responses upon up-regulation of *PRPS1* and *PRPL4* gene expression and the consequent alteration of plastid protein homeostasis. Whereas the *PRPS1*-related response mainly relies on chloroplast breakdown with degradation and loss of chloroplast proteins, nucleic acids, pigments, lipids, and polysaccharides, aimed to promote the remobilization of resources, such as carbon and nitrogen, in favour of the anticipated reproductive phase, i.e. an escape strategy from a stress condition, the *PRPL4*-related response consists in the activation of a stress pathway, compatible with the chloroplast UPR.

## Author Contributions

L.T. and P.P. designed the study. N.J., S.F., and L.T. took care of the isolation and the phenotypical characterization of mutants and transgenic lines. L.T. and N.J. performed the molecular characterization of plant lines and bacterial strains. S.M. and N.J. carried out the characterization of embryo-lethal mutants. A.C., N.J. and F.Z. performed analyses of transcriptomic data. G.D., E.C., M.M. and C.V. performed proteomic analyses. L.T., N.J., F.B. and P.P. drafted the manuscript. L.T., N.J. and P.P. coordinated the study and took care of the final version of the manuscript. All authors have read and agreed to the published version of the manuscript.

## Data Availability

The data that support the findings of this study are openly available in Gene Expression Omnibus at https://www.ncbi.nlm.nih.gov/geo/, reference number GSE205271, and in ProteomeXchange Consortium at http://www.proteomexchange.org/, reference number PXD034479.

## Acknowledgments

We are grateful to James Friel for critical reading of the manuscript and English editing. We are thankful to Lucio Conti and Cecilia Zumajo for fruitful discussion on flowering time regulatory mechanisms. We are also grateful to Norma Lattuada, Roberto Ferrari, Valerio Paravicini and Mario Beretta for excellent technical assistance. NoLimits platform at University of Milano is acknowledged for TEM analyses.

## References

Ahmed T, Yin Z, Bhushan S. 2016. Cryo-EM structure of the large subunit of the spinach chloroplast ribosome. Scientific Reports 6: 1–13.

Arcus V. 2002. OB-fold domains: A snapshot of the evolution of sequence, structure and function. Current Opinion in Structural Biology 12: 794–801.

Asakura Y, Barkan A. 2006. Arabidopsis orthologs of maize chloroplast splicing factors promote splicing of orthologous and species-specific group II introns. Plant Physiology 142: 1656–1663.

Barkan A. 1998. Approaches to investigating nuclear genes that function in chloroplast biogenesis in land plants. Methods in Enzymology 297: 38–57.

Beckert B, Turk M, Czech A, Berninghausen O, Beckmann R, Ignatova Z, Plitzko JM, Wilson DN. 2018. Structure of a hibernating 100S ribosome reveals an inactive conformation of the ribosomal protein S1. Nature Microbiology 3: 1115–1121.

Bieri P, Leibundgut M, Saurer M, Boehringer D, Ban N. 2017. The complete structure of the chloroplast 70S ribosome in complex with translation factor pY. The EMBO Journal 36: 475–486.

Bolouri Moghaddam MR, Van den Ende W. 2013. Sugars, the clock and transition to flowering. Frontiers in Plant Science 4.

Boni I V., Artamonova VS, Dreyfus M. 2000. The last RNA-binding repeat of the Escherichia coli ribosomal protein S1 is specifically involved in autogenous control. Journal of Bacteriology 182: 5872–5879.

Briani F, Curti S, Rossi F, Carzaniga T, Mauri P, Dehò G. 2008. Polynucleotide phosphorylase hinders mRNA degradation upon ribosomal protein S1 overexpression in Escherichia coli. RNA 14: 2417–2429.

Bryant N, Lloyd J, Sweeney C, Myouga F, Meinke D. 2011. Identification of nuclear genes encoding chloroplast-localized proteins required for embryo development in Arabidopsis. Plant Physiology 155: 1678–1689.

Bubunenko MG, Schmidt J, Subramanian AR. 1994. Protein substitution in chloroplast ribosome evolution a eukaryotic cytosolic protein has replaced its organelle homologue (L23) in spinach. Journal of Molecular Biology 240: 28–41.

Bycroft M, Hubbard TJP, Proctor M, Freund SMV, Murzin AG. 1997. The solution structure of the S1 RNA binding domain: a member of an ancient nucleic acid-binding fold. Cell 88: 235–242.

Byrgazov K, Grishkovskaya I, Arenz S, Coudevylle N, Temmel H, Wilson DN, Djinovic-Carugo K, Moll I. 2015. Structural basis for the interaction of protein S1 with the Escherichia coli ribosome. Nucleic Acids Research 43: 661–673.

Chalupska D, Lee HY, Faris JD, Evrard A, Chalhoub B, Haselkorn R, Gornicki P. 2008. Acc homoeoloci and the evolution of wheat genomes. Proceedings of the National Academy of Sciences of the United States of America 105: 9691–9696.

Chen H, Hwang JE, Lim CJ, Kim DY, Lee SY, Lim CO. 2010. Arabidopsis DREB2C functions as a transcriptional activator of HsfA3 during the heat stress response. Biochemical and Biophysical Research Communications 401: 238–244.

Cifuentes-Goches JC, Hernández-Ancheyta L, Guarneros G, Oviedo N, Hernández-Sánchez J. 2019. Domains two and three of Escherichia coli ribosomal S1 protein confers 30S subunits a high affinity for downstream A/U-rich mRNAs. Journal of Biochemistry 166: 29–40.

Colombo M, Tadini L, Peracchio C, Ferrari R, Pesaresi P. 2016. GUN1, a Jack-Of-All-Trades in Chloroplast Protein Homeostasis and Signaling. Frontiers in Plant Science 7.

Czechowski T, Stitt M, Altmann T, Udvardi MK, Scheible WR. 2005. Genome-wide identification and testing of superior reference genes for transcript normalization in arabidopsis. Plant Physiology 139: 5–17.

Delvillani F, Papiani G, Dehó G, Briani F. 2011. S1 ribosomal protein and the interplay between translation and mRNA decay. Nucleic Acids Research 39: 7702–7715.

Duval M, Korepanov A, Fuchsbauer O, Fechter P, Haller A, Fabbretti A, Choulier L, Micura R, Klaholz BP, Romby P, et al. 2013. Escherichia coli Ribosomal Protein S1 Unfolds Structured mRNAs Onto the Ribosome for Active Translation Initiation. PLoS Biology 11: e1001731.

Fernández-Bautista N, Fernández-Calvino L, Muñoz A, Castellano MM. 2017. HOP3, a member of the HOP family in Arabidopsis, interacts with BiP and plays a major role in the ER stress response. Plant Cell and Environment 40: 1341–1355.

Franzetti B, Carol P, Maches R. 1992. Characterization and RNA-binding properties of a chloroplast S1-like ribosomal protein. Journal of Biological Chemistry 267: 19075–19081.

Freytes SN, Canelo M, Cerdán PD. 2021. Regulation of Flowering Time: When and Where? Current Opinion in Plant Biology 63.

Giorginis S, Subramanian AR. 1980. The major ribosome binding site of Escherichia coli ribosomal protein S1 is located in its N-terminal segment. Journal of Molecular Biology 141: 393–408.

Graf M, Arenz S, Huter P, Dönhöfer A, Nováček J, Wilson DN. 2017. Cryo-EM structure of the spinach chloroplast ribosome reveals the location of plastid-specific ribosomal proteins and extensions. Nucleic Acids Research 45: 2887–2896.

Hajnsdorf E, Boni I V. 2012. Multiple activities of RNA-binding proteins S1 and Hfq. Biochimie 94: 1544–1553.

Hess WR, Hoch B, Zeltz P, Hübschmann T, Kössel H, Börner T. 1994. Inefficient rpl2 splicing in barley mutants with ribosome-deficient plastids. Plant Cell 6: 1455–1465.

Hirose T, Sugiura M. 2004. Multiple elements required for translation of plastid atpB mRNA lacking the Shine-Dalgarno sequence. Nucleic Acids Research 32: 3503–3510.

Hsieh HL, Okamoto H, Wang M, Ang LH, Matsui M, Goodman H, Deng XW. 2000. FIN219, an auxin-regulated gene, defines a link between phytochrome A and the downstream regulator COP1 in light control of Arabidopsis development. Genes and Development 14: 1958–1970.

Jeran N, Rotasperti L, Frabetti G, Calabritto A, Pesaresi P, Tadini L. 2021. The PUB4 E3 ubiquitin ligase is responsible for the variegated phenotype observed upon alteration of chloroplast protein homeostasis in arabidopsis cotyledons. Genes 12: 1387.

Kalapos MP, Paulus H, Sarkar N. 1997. Identification of ribosomal protein S1 as a poly(A) binding protein in Escherichia coli. Biochimie 79: 493–502.

Karim S, Holmström KO, Mandal A, Dahl P, Hohmann S, Brader G, Palva ET, Pirhonen M. 2007. AtPTR3, a wound-induced peptide transporter needed for defence against virulent bacterial pathogens in Arabidopsis. Planta 225: 1431–1445.

Kazan K, Lyons R. 2016. The link between flowering time and stress tolerance. Journal of Experimental Botany 67: 47–60.

Kitakawa M, Isono K. 1982. An amber mutation in the gene rpsA for ribosomal protein S1 in Escherichia coli. Molecular and General Genetics MGG 185: 445–447.

Kmiec B, Teixeira PF, Glaser E. 2014. Shredding the signal: Targeting peptide degradation in mitochondria and chloroplasts. Trends in Plant Science 19: 771–778.

Lakhssassi N, Doblas VG, Rosado A, del Valle AE, Posé D, Jimenez AJ, Castillo AG, Valpuesta V, Borsani O, Botella MA. 2012. The Arabidopsis TETRATRICOPEPTIDE THIOREDOXIN- LIKE gene family is required for osmotic stress tolerance and male sporogenesis. Plant Physiology 158: 1252–1266.

Lemke MD, Fisher KE, Kozlowska MA, Tano DW, Woodson JD. 2021. The core autophagy machinery is not required for chloroplast singlet oxygen-mediated cell death in the Arabidopsis thaliana plastid ferrochelatase two mutant. BMC Plant Biology 21: 1–20.

Li B, Dewey CN. 2011. RSEM: Accurate transcript quantification from RNA-Seq data with or without a reference genome. BMC Bioinformatics 12: 1–16.

Li S, Fu Q, Chen L, Huang W, Yu D. 2011. Arabidopsis thaliana WRKY25, WRKY26, and WRKY33 coordinate induction of plant thermotolerance. Planta 233: 1237–1252.

Liu W, Liu Z, Mo Z, Guo S, Liu Y, Xie Q. 2021. ATG8-Interacting Motif: Evolution and Function in Selective Autophagy of Targeting Biological Processes. Frontiers in Plant Science 12: 2671.

Llamas E, Pulido P, Rodriguez-Concepcion M. 2017. Interference with plastome gene expression and Clp protease activity in Arabidopsis triggers a chloroplast unfolded protein response to restore protein homeostasis. PLoS Genetics 13: 1–28.

Mache R. 1990. Chloroplast ribosomal proteins and their genes. Plant Science 72: 1–12.

Merendino L, Falciatore A, Rochaix JD. 2003. Expression and RNA binding properties of the chloroplast ribosomal protein S1 from Chlamydomonas reinhardtii. Plant Molecular Biology 53: 371–382.

Mi H, Ebert D, Muruganujan A, Mills C, Albou LP, Mushayamaha T, Thomas PD. 2021. PANTHER version 16: A revised family classification, tree-based classification tool, enhancer regions and extensive API. Nucleic Acids Research 49: D394–D403.

Michaeli S, Honig A, Levanony H, Peled-Zehavi H, Galili G. 2014. Arabidopsis ATG8- INTERACTING PROTEIN1 is involved in autophagy-dependent vesicular trafficking of plastid proteins to the vacuole. The Plant Cell 26: 4084–4101.

Mishra RC, Grover A. 2016. ClpB/Hsp100 proteins and heat stress tolerance in plants. Critical Reviews in Biotechnology 36: 862–874.

Murzin AG. 1993. OB(oligonucleotide/oligosaccharide binding)-fold: Common structural and functional solution for non-homologous sequences. EMBO Journal 12: 861–867.

Nagashima Y, Ohshiro K, Iwase A, Nakata MT, Maekawa S, Horiguchi G. 2020. The bRPS6- family protein RFC3 prevents interference by the splicing factor CFM3b during plastid rRNA biogenesis in Arabidopsis thaliana. Plants 9.

Nishizawa A, Yabuta Y, Shigeoka S. 2008. Galactinol and raffinose constitute a novel function to protect plants from oxidative damage. Plant Physiology 147: 1251–1263.

Nishizawa A, Yabuta Y, Yoshida E, Maruta T, Yoshimura K, Shigeoka S. 2006. Arabidopsis heat shock transcription factor A2 as a key regulator in response to several types of environmental stress. Plant Journal 48: 535–547.

Ostheimer GJ, Williams-Carrier R, Belcher S, Osborne E, Gierke J, Barkan A. 2003. Group II intron splicing factors derived by diversification of an ancient RNA-binding domain. EMBO Journal 22: 3919–3929.

Paieri F, Tadini L, Manavski N, Kleine T, Ferrari R, Morandini P, Pesaresi P, Meurer J, Leister D. 2018. The DEAD-box RNA helicase RH50 is a 23S-4.5S rRNA maturation factor that functionally overlaps with the plastid signaling factor GUN1. Plant Physiology 176: 634–648.

Panikulangara TJ, Eggers-Schumacher G, Wunderlich M, Stransky H, Schöffl F. 2004. Galactinol synthase1. A novel heat shock factor target gene responsible for heat-induced synthesis of raffinose family oligosaccharides in arabidopsis. Plant Physiology 136: 3148–3158.

Paradiso A, Domingo G, Blanco E, Buscaglia A, Fortunato S, Marsoni M, Scarcia P, Caretto S, Vannini C, de Pinto MC. 2020. Cyclic AMP mediates heat stress response by the control of redox homeostasis and ubiquitin-proteasome system. Plant Cell and Environment 43: 2727–2742.

Pérez-Martín M, Pérez-Pérez ME, Lemaire SD, Crespo JL. 2014. Oxidative stress contributes to autophagy induction in response to endoplasmic reticulum stress in chlamydomonas reinhardtii. Plant Physiology 166: 997–1008.

Perez-Riverol Y, Csordas A, Bai J, Bernal-Llinares M, Hewapathirana S, Kundu DJ, Inuganti A, Griss J, Mayer G, Eisenacher M, et al. 2019. The PRIDE database and related tools and resources in 2019: Improving support for quantification data. Nucleic Acids Research 47: D442– D450.

Pesaresi P, Varotto C, Meurer J, Jahns P, Salamini F, Leister D. 2001. Knock-out of the plastid ribosomal protein L11 in Arabidopsis: Effects on mRNA translation and photosynthesis. The Plant Journal 27: 179–189.

Phee BK, Kim J Il, Shin DH, Yoo J, Park KJ, Han YJ, Kwon YK, Cho MH, Jeon JS, Bhoo SH, et al. 2008. A novel protein phosphatase indirectly regulates phytochrome-interacting factor 3 via phytochrome. Biochemical Journal 415: 247–255.

Pulido P, Llamas E, Llorente B, Ventura S, Wright LP, Rodríguez-Concepción M. 2016. Specific Hsp100 Chaperones Determine the Fate of the First Enzyme of the Plastidial Isoprenoid Pathway for Either Refolding or Degradation by the Stromal Clp Protease in Arabidopsis. PLoS Genetics 12.

Ramundo S, Casero D, Muhlhaus T, Hemme D, Sommer F, Crevecoeur M, Rahire M, Schroda M, Rusch J, Goodenough U, et al. 2014. Conditional Depletion of the Chlamydomonas Chloroplast ClpP Protease Activates Nuclear Genes Involved in Autophagy and Plastid Protein Quality Control. The Plant Cell 26: 2201–2222.

Ramundo S, Rochaix JD. 2014. Chloroplast unfolded protein response, a new plastid stress signaling pathway? Plant Signaling and Behavior 9: 1–3.

Robinson MD, McCarthy DJ, Smyth GK. 2009. edgeR: A Bioconductor package for differential expression analysis of digital gene expression data. Bioinformatics 26: 139–140.

Romani I, Tadini L, Rossi F, Masiero S, Pribil M, Jahns P, Kater M, Leister D, Pesaresi P. 2012. Versatile roles of Arabidopsis plastid ribosomal proteins in plant growth and development. Plant Journal 72: 922–934.

Rottet S, Besagni C, Kessler F. 2015. The role of plastoglobules in thylakoid lipid remodeling during plant development. Biochimica et Biophysica Acta - Bioenergetics 1847: 889–899.

Sakuma Y, Maruyama K, Qin F, Osakabe Y, Shinozaki K, Yamaguchi-Shinozaki K. 2006. Dual function of an Arabidopsis transcription factor DREB2A in water-stress-responsive and heat-stress- responsive gene expression. Proceedings of the National Academy of Sciences of the United States of America 103: 18822–18827.

Salah P, Bisaglia M, Aliprandi P, Uzan M, Sizun C, Bontems F. 2009. Probing the relationship between gram-negative and gram-positive S1 proteins by sequence analysis. Nucleic Acids Research 37: 5578–5588.

Samalova M, Brzobohaty B, Moore I. 2005. pOp6/LhGR: a stringently regulated and highly responsive dexamethasone-inducible gene expression system for tobacco. The Plant Journal 41: 919– 935.

Sasaki I, Bertani G. 1965. Growth abnormalities in Hfr derivatives of Escherichia coli strain C. Journal of general microbiology 40: 365–376.

Sawada Y, Toyooka K, Kuwahara A, Sakata A, Nagano M, Saito K, Hirai MY. 2009. Arabidopsis bile acid:sodium symporter family protein 5 is involved in methionine-derived glucosinolate biosynthesis. Plant and Cell Physiology cell physiology 50: 1579–1586.

Schägger H, von Jagow G. 1987. Tricine-sodium dodecyl sulfate-polyacrylamide gel electrophoresis for the separation of proteins in the range from 1 to 100 kDa. Analytical Biochemistry 166: 368–379.

Schulte W, Töpfer R, Stracke R, Schell J, Martini N. 1997. Multi-functional acetyl-CoA carboxylase from Brassica napus is encoded by a multi-gene family: Indication for plastidic localization of at least one isoform. Proceedings of the National Academy of Sciences of the United States of America 94: 3465–3470.

Schuster J, Knill T, Reichelt M, Gershenzon J, Binder S. 2006. BRANCHED-CHAIN AMINOTRANSFERASE4 is part of the chain elongation pathway in the biosynthesis of methionine- derived glucosinolates in Arabidopsis. The Plant Cell 18: 2664–2679.

Sjögren LLE, MacDonald TM, Sutinen S, Clarke AK. 2004. Inactivation of the clpC1 gene encoding a chloroplast Hsp100 molecular chaperone causes growth retardation, leaf chlorosis, lower photosynthetic activity, and a specific reduction in photosystem content. Plant Physiology 136: 4114– 4126.

Skouv J, Schnier J, Rasmussen MD, Subramanian AR, Pedersen S. 1990. Ribosomal protein S1 of Escherichia coli is the effector for the regulation of its own synthesis. Journal of Biological Chemistry 265: 17044–17049.

Sørensen MA, Fricke J, Pedersen S. 1998. Ribosomal protein S1 is required for translation of most, it not all, natural mRNAs in Escherichia coli in vivo. Journal of Molecular Biology 280: 561–569.

Subramanian AR. 1983. Structure and Functions of Ribosomal Protein S1. Progress in Nucleic Acid Research and Molecular Biology 28: 101–142.

Sugita M, Sugita C, Sugiura M. 1995. Structure and expression of the gene encoding ribosomal protein S1 from the cyanobacterium Synechococcus sp. strain PCC 6301: striking sequence similarity to the chloroplast ribosomal protein CS1. Molecular and General Genetics MGG 246: 142–147.

Sukhodolets M V., Garges S, Adhya S. 2006. Ribosomal protein S1 promotes transcriptional cycling. RNA 12: 1505–1513.

Sun W, Bernard C, Van Cotte B De, Van Montagu M, Verbruggen N. 2001. At-HSP17.6A, encoding a small heat-shock protein in Arabidopsis, can enhance osmotolerance upon overexpression. The Plant Journal 27: 407–415.

Supek F, Bošnjak M, Škunca N, Šmuc T. 2011. Revigo summarizes and visualizes long lists of gene ontology terms. PLoS ONE 6: e21800.

Tadini L, Ferrari R, Lehniger M-K, Mizzotti C, Moratti F, Resentini F, Colombo M, Costa A, Masiero S, Pesaresi P. 2018. Trans-splicing of plastid rps12 transcripts, mediated by AtPPR4, is essential for embryo patterning in Arabidopsis thaliana. Planta 248: 257–265.

Tadini L, Jeran N, Peracchio C, Masiero S, Colombo M, Pesaresi P. 2020a. The plastid transcription machinery and its coordination with the expression of nuclear genome: Plastid-Encoded Polymerase, Nuclear-Encoded Polymerase and the Genomes Uncoupled 1-mediated retrograde communication. Philosophical Transactions of the Royal Society B: Biological Sciences 375: 20190399.

Tadini L, Jeran N, Pesaresi P. 2020b. GUN1 and Plastid RNA Metabolism: Learning from Genetics. Cells 9: 2307.

Tadini L, Peracchio C, Trotta A, Colombo M, Mancini I, Jeran N, Costa A, Faoro F, Marsoni M, Vannini C, et al. 2020c. GUN1 influences the accumulation of NEP-dependent transcripts and chloroplast protein import in Arabidopsis cotyledons upon perturbation of chloroplast protein homeostasis. The Plant Journal 101: 1198–1220.

Tadini L, Pesaresi P, Kleine T, Rossi F, Guljamow A, Sommer F, Mühlhaus T, Schroda M, Masiero S, Pribil M, et al. 2016. GUN1 Controls Accumulation of the Plastid Ribosomal Protein S1 at the Protein Level and Interacts with Proteins Involved in Plastid Protein Homeostasis. Plant physiology 170: 1817–30.

Tadini L, Romani I, Pribil M, Jahns P, Leister D, Pesaresi P. 2012. Thylakoid redox signals are integrated into organellar-gene-expression-dependent retrograde signaling in the prors1-1 mutant. Frontiers in Plant Science 3: 1–13.

Takeno K. 2016. Stress-induced flowering: The third category of flowering response. Journal of Experimental Botany 67: 4925–4934.

Theobald DL, Mitton-Fry RM, Wuttke DS. 2003. Nucleic acid recognition by OB-fold proteins. Annual Review of Biophysics and Biomolecular Structure 32: 115–133.

Tian T, Liu Y, Yan H, You Q, Yi X, Du Z, Xu W, Su Z. 2017. AgriGO v2.0: A GO analysis toolkit for the agricultural community, 2017 update. Nucleic Acids Research 45: W122–W129.

Tiller N, Bock R. 2014. The translational apparatus of plastids and its role in plant development. Molecular Plant 7: 1105–1120.

Toribio R, Mangano S, Fernández-Bautista N, Muñoz A, Castellano MM. 2020. HOP, a Co- chaperone Involved in Response to Stress in Plants. Frontiers in Plant Science 11: 1657.

Tran LSP, Nakashima K, Sakuma Y, Simpson SD, Fujita Y, Maruyama K, Fujita M, Seki M, Shinozaki K, Yamaguchi-Shinozaki K. 2004. Isolation and functional analysis of arabidopsis stress-inducible NAC transcription factors that bind to a drought-responsive cis-element in the early responsive to dehydration stress 1 promoter. The Plant Cell 16: 2481–2498.

Vannini C, Domingo G, Fiorilli V, Seco DG, Novero M, Marsoni M, Wisniewski-Dye F, Bracale M, Moulin L, Bonfante P. 2021. Proteomic analysis reveals how pairing of a Mycorrhizal fungus with plant growth-promoting bacteria modulates growth and defense in wheat. Plant Cell and Environment 44: 1946–1960.

van Wijk KJ. 2015. Protein Maturation and Proteolysis in Plant Plastids, Mitochondria, and Peroxisomes. Annual Review of Plant Biology 66: 75–111.

Van Wijk KJ, Kessler F. 2017. Plastoglobuli: Plastid Microcompartments with Integrated Functions in Metabolism, Plastid Developmental Transitions, and Environmental Adaptation. Annual Review of Plant Biology 68: 253–289.

Wiśniewski JR. 2019. Filter Aided Sample Preparation A tutorial. Analytica Chimica Acta 1090: 23–30.

Woodson JD. 2016. Chloroplast quality control - balancing energy production and stress. The New phytologist 212: 36–41.

Woodson JD. 2022. Control of chloroplast degradation and cell death in response to stress. Trends in Biochemical Sciences.

Wu GZ, Meyer EH, Richter AS, Schuster M, Ling Q, Schöttler MA, Walther D, Zoschke R, Grimm B, Jarvis RP, et al. 2019. Control of retrograde signalling by protein import and cytosolic folding stress. Nature Plants 5: 525–538.

Yamaguchi K, Von Knoblauch K, Subramanian AR. 2000. The plastid ribosomal proteins. IDENTIFICATION OF ALL THE PROTEINS IN THE 30 S SUBUNIT OF AN ORGANELLE RIBOSOME (CHLOROPLAST)*. Journal of Biological Chemistry 275: 28455–28465.

Yamaguchi K, Subramanian AR. 2003. Proteomic identification of all plastid-specific ribosomal proteins in higher plant chloroplast 30S ribosomal subunit. European journal of biochemistry 270: 190–205.

Yin T, Pan G, Liu H, Wu J, Li Y, Zhao Z, Fu T, Zhou Y. 2012. The chloroplast ribosomal protein L21 gene is essential for plastid development and embryogenesis in Arabidopsis. Planta 235: 907– 921.

Yu HD, Yang XF, Chen ST, Wang YT, Li JK, Shen Q, Liu XL, Guo FQ. 2012. Downregulation of chloroplast RPS1 negatively modulates nuclear heat-responsive expression of HsfA2 and its target genes in Arabidopsis. PLoS Genetics 8.

Zechmann B. 2019. Ultrastructure of plastids serves as reliable abiotic and biotic stress marker. PLoS ONE 14.

Zhou K, Zhang C, Xia J, Yun P, Wang Y, Ma T, Li Z. 2021. Albino seedling lethality 4; Chloroplast 30S Ribosomal Protein S1 is Required for Chloroplast Ribosome Biogenesis and Early Chloroplast Development in Rice. Rice 14: 1–12.

Zhuang X, Jiang L. 2019. Chloroplast degradation: Multiple routes into the vacuole. Frontiers in Plant Science 10: 359.

Zoschke R, Bock R. 2018. Chloroplast translation: Structural and functional organization, operational control, and regulation. Plant Cell 30: 745–770.

Zubko MK, Day A. 1998. Stable albinism induced without mutagenesis: A model for ribosome-free plastid inheritance. The Plant Journal 15: 265–271.

